# Sensory neurons shape macrophage identity via TGF-β signalling

**DOI:** 10.1101/2025.02.06.635770

**Authors:** Julia Kolter, Clarissa-Laura Döring, Sebastian Baasch, Reem Alsumati, Zohreh Mansoori Moghadam, Vitka Gres, Florens Lohrmann, Gabriel Victor Lucena da Silva, Conceicao Elidianne Anibal Silva, Theresa Buchegger, Layal Doumard, Benjamin Voisin, Nico Lachmann, Katrin Kierdorf, Vincent Flacher, Philipp Henneke

## Abstract

Macrophages play integral roles in maintaining homeostasis and function in their tissues of residence. In the skin, prenatally seeded and highly specialized macrophages physically interact with sensory nerves and contribute to their regeneration after injury. However, mechanisms underlying the development and maintenance of this potentially lifelong commitment of macrophages to nociceptors remain largely elusive. Here, we found that infiltrating myeloid progenitor cells approached the sprouting axons of sensory nerves and gradually adopted a nerve-associated macrophage-like profile. This change in identity was steered and maintained by the immediate microenvironment, in particular TGF-β, which was locally activated by the physical interaction with nerves and integrin-mediated cleavage. Following injury, TGF-β driven specification of macrophages essentially supported nerve regeneration. Overall, we identified TGF-β as a central mediator governing local imprinting and long-term specialization of macrophages in the skin, providing insights into the bidirectional communication between macrophages and sensory nerves.

## INTRODUCTION

The skin is a large interface organ of the human body and acts as a critical protective barrier to the environment. As in most tissues, macrophages represent the most abundant resident immune cell type in the skin with important roles in tissue homeostasis, barrier defense and inflammation. Macrophages are able to adapt to different microenvironments and perform a diversity of tissue specific functions. Consequently, macrophages of different organs display significant variations in immunophenotype, morphology, transcriptome, and function, underscoring their distinct roles in maintaining tissue homeostasis. In general, tissue-specific cues, such as metabolites, growth factors and microbial products, are thought to contribute to the induction of distinct transcriptional programs (Mass et al., 2023). However, factors directing precise diversification in a structurally complex tissue like the skin are poorly defined. In contrast, some insights have been gained from situations, when the cellular macrophage environment is less complex or better defined. Alveolar macrophages, for instance, are exposed to Granulocyte-macrophage colony-stimulating factor (GM-CSF) and transforming growth factor (TGF)-β in the alveolar spaces, resulting in activation of the master transcription factor peroxisome proliferator-activated receptor (PPAR) -γ, which drives expression of signature gene and thus differentiation early in life (Schneider et al., 2014; Yu et al., 2017). Large peritoneal macrophages induce the transcription factor GATA6 due to the presence of the vitamin A metabolite retinoic acid in the visceral body cavities (Buechler et al., 2019; Okabe and Medzhitov, 2014). In extension of these models, tissue complexity with diverse anatomical niches requires the acquisition of discrete functions by macrophage types residing within the same organ, for instance in the meninges and choroid plexus of the brain or in the red pulp and marginal zones of the spleen. Moreover, at an even higher resolution, highly specialized macrophages were found to associate to substructures within tissues, such as blood vessels and peripheral nerves (Abtin et al., 2014; Chakarov et al., 2019). We previously reported the existence of a small subset of dermal macrophages, which is associated with sensory nerves, patrols axons by migration and contributes to nerve repair upon injury (Kolter et al., 2019). These sensory nerve-associated macrophages (sNAM) exhibit a distinct transcriptional profile and immunophenotype in the dermis, characterized e.g., by high expression of the fractalkine receptor CX3CR1 and low expression of the mannose receptor CD206. Interestingly, sNAM are of embryonic origin and capable of self-maintenance, unlike the majority of interstitial dermal macrophages, which are rapidly replaced by circulating monocytes in steady state (Kolter et al., 2019; Tamoutounour et al., 2013). These findings are in line with the generally accepted model that the origin of tissue macrophages is multifaceted with tissue-specific contributions from both embryonic and adult hematopoiesis, ranging from exclusively yolk-sac derived microglia in the adult central nervous system (CNS) to mostly blood monocyte-derived macrophages in the intestinal wall (Bain et al., 2014; Ginhoux et al., 2010; Mass et al., 2023; Schulz et al., 2012). Nevertheless, even in the gut, self-renewing macrophages are found in vicinity of specific structures including enteric neurons and submucosal vasculature (De Schepper et al., 2018), suggesting that tissue niche signals instruct longevity. In comparison to the situation in the CNS, macrophages of the peripheral nervous system (PNS) remain incompletely understood (Kolter et al., 2020). PNS macrophages within the dorsal root ganglia (DRG) and larger nerves, such as the vagal and the sciatic nerves, were shown to be of mixed origin and to overlap in their transcriptional signature with activated microglia (Wang et al., 2020). Yet, PNS macrophages within the larger nerves are heterogeneous, comprising distinct epineurial and endoneurial macrophages (Ydens et al., 2020). In the enteric nervous system, muscularis macrophages in the myenteric plexus coordinate gastrointestinal mobility and support neuronal survival and maturation early in life (De Schepper et al., 2018; Muller et al., 2020; Viola et al., 2023). However, in the gut and most other tissues, it is challenging to distinguish between nerve-associated macrophages and interstitial macrophages due to overlapping marker expression and lack of physical separation, hence their origin and discriminative functions remain elusive. In addition, the low cellular number of this rare cell type requires highly dedicated toolboxes. Thus, the molecular determinants of specific nerve-associated macrophages, e.g., the cues provided by different types of peripheral nerves, remain to be established.

Here, we combined *in vitro* and *in vivo* approaches to identify the molecular signals and their sources which mediate the unique interaction between macrophages and sensory nerves in the dermis. We found macrophages to be rapidly recruited to sprouting axons during development and after injury, which was accompanied with profound transcriptional and phenotypic changes. Transcriptional profiling revealed a critical role for TGF-β in the differentiation of sNAM, which was locally activated by integrin-expressing macrophages, and contributed to sNAM self-maintenance as well as recruitment and differentiation of new sNAM-like macrophages after injury. Accordingly, disruption of the TGF-β-mediated crosstalk between neurons and macrophages delayed peripheral nerve regeneration after physical injury, underscoring the importance of specialized immune cells in reinstating tissue homeostasis.

## RESULTS

### sNAM recruited upon injury derive from bone marrow progenitors

Upon injury, sensory nerves sprout and form a dense interconnected axon network at the site of injury, a so-called regeneration ring (Enamorado et al., 2023; Kolter et al., 2019). Adult sNAM are of prenatal origin and self-maintain in steady state. However, we found that macrophages colonizing the newly sprouted nerves in the regeneration ring were not derived from sNAM residing on nerves before the injury, even though they exhibited the characteristic sNAM immunophenotype, e.g., high CX_3_CR1 and low CD206 expression (Kolter et al., 2019). Since these newly formed sNAM also appeared in the absence of circulating monocytes, we concluded that interstitial macrophages could transform into sNAM. Here, we aimed to rigorously explore the stepwise adaptation process to the neuronal niche with novel fate mapping approaches. First, we exploited that sNAM are unique among dermal macrophages with respect to robust CX3CR1 expression. Thus, we induced *Cx3cr1^creERT2/gfp^ R26^tdT/tdT^* mice with tamoxifen (TAM) before injury to label *bona fide* sNAM in the dermis. *Bona fide*, prior resident sNAM (i.e., macrophages associated with β3-tubulin^+^ sensory nerves outside of the injured area) were reliably labelled with GFP and tdTomato. In contrast, sNAM-like macrophages associated with axons in the regeneration ring only expressed GFP, indicating that these cells did not originate from the prior resident sNAM (Fig. 1A). In line with their acquisition of CX3CR1, macrophages in the regeneration ring expressed tdTomato if tamoxifen was administered after the injury (Fig. S1A, B). Next, we assessed whether these newly recruited cells were indeed derived from differentiated interstitial macrophages. With the exception of sNAM, all dermal macrophages strongly express the mannose receptor CD206 (Kolter et al., 2019). Hence, we employed the recently developed *Mrc1^creERT2/creERT2^ R26^tdT/tdT^* mice, in which more than 90% of CD206-expressing dermal macrophages were labelled with tdTomato after TAM induction (Forde et al., 2023; Masuda et al., 2022). In contrast, blood monocytes were tdTomato-negative (Fig. S1C). Secondly, we used *Cxcr4^creERT2/+^ R26^tdT/tdT^* mice, in which cells derived from hematopoietic stem cells expressed tdTomato (Werner et al., 2020) (Fig. S1D). The combination of these two approaches enabled us to track the origin of the majority of dermal macrophages. Upon punch injury, most interstitial macrophages remained labelled in the *Mrc1^creERT2/creERT2^ R26^tdT/tdT^* mice, whereas recruited sNAM in the regeneration ring were negative for tdTomato (Fig. 1B-D). Instead, they showed strong labelling in *Cxcr4^creERT2/+^R26^tdT/tdT^* mice (Fig. 1B, E, F). These findings indicated that these sNAM did not derive from prior tissue-resident and fully differentiated interstitial dermal macrophages, but rather from HSC-derived progenitors which migrated into the injury site and gave rise to sNAM. It seems worth noting that endothelial cells also highly expressed tdTomato in *Cxcr4^creERT2/+^R26^tdT/tdT^*mice, which hampered the immunofluorescence analysis (Fig. S1E, F). To complement the analysis, we hence performed bone marrow transplantation in ear-shielded mice 8 weeks before injury (Fig. 1G and S1G-I). In agreement with the data generated in TAM-induced reporter mice, ear-shielded *Cd45.1* mice transplanted with *Cx3cr1^gfp/+^* bone marrow showed robust recruitment of GFP^+^ CX3CR1^high^ cells into the regeneration ring, while the recruited cells in *Cx3cr1^gfp/+^* mice transplanted with *Cd45.1* bone marrow did not express GFP (Fig. 1G-I and Fig. S1J, K). These results strongly argued that sNAM recruited after injury derived from hematopoietic stem cells and not from resident interstitial dermal macrophages. However, when inducing *Mrc1^creERT2/creERT2^ R26^tdT/tdT^* mice simultaneously with the punch injury, many sNAM within the regeneration ring were labeled 14 days post injury, while monocytes did not express tdTomato at any stage (Fig. 1J, S1L). This finding is in agreement with our hypothesis that acquisition of the unique sNAM phenotype is a stepwise process, where monocytes enter the dermis and upregulate CD206 before concluding their adaptation process to become sNAM. However, these data add another layer of complexity, in that phenotypically mature interstitial macrophages only transiently keep flexibility for further specification. Nevertheless, the dynamic frame of this process remained to be explored.

**Figure 1.**
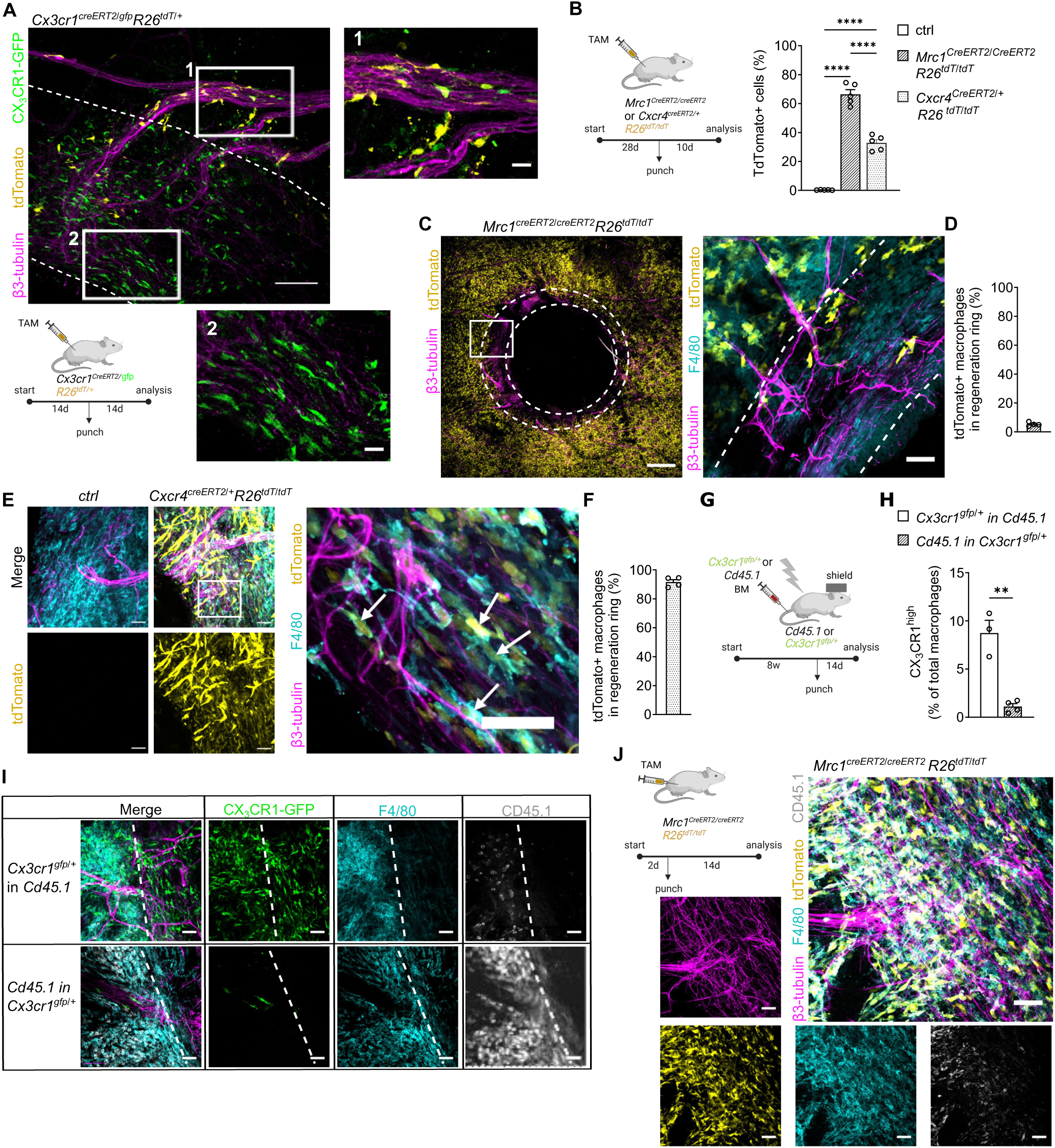
sNAM recruited upon injury derive from bone marrow progenitors. **(A)** Experimental setup and images showing lack of tdTomato labelling in the regeneration ring of ear punches of *Cx3cr1creERT2/gfp R26tdT/tdT* mice, induced prior to injury. Dashed lines indicate regeneration rings at injury center (n=4). Scale bar: 100μm. **(B)** Experimental setup and tdTomato expression in all CD64+ dermal macrophages in *Mrc1creERT2/creERT2* or *Cxcr4creERT2/+ R26tdT/tdT* 10 days post injury, measured via flow cytometry (n=5 mice per group). **(C)** Image depicting absence of tdTomato expression in the regeneration ring (indicated by dotted lines) of ear punches of *Mrc1creERT2/creERT2 R26tdT/tdT*mice in (C). Scale bar: 500μm, zoom-in image 50μm. **(D)** Quantification of tdTomato+ F4/80+ cells in regeneration rings (D). **(E)** Image depicting tdTomato expression in macrophages in ear punches of *Cxcr4creERT2/+ R26tdT/tdT* mice in (C). Scale bar: 50μm, zoom-in image 40μm. Arrows point to exemplary TdTomato+ nerve-associated macrophages. **(F)** Quantification of tdTomato+ F4/80+ cells in regeneration rings (F). **(G)** Ear-shielded, irradiated CD45.1 mice were transplanted with *Cx3cr1gfp/+* and vice versa, punched 8 weeks post transfer and analyzed 14 days later. **(H)** Quantification of CX3CR1high macrophages in chimeric mice 14 days post punch injury (n=3-4 per group) (two-tailed unpaired *t*-test). **(I)** Whole mount images depicting CX3CR1high macrophages in ear punches in mice in which either donor (upper panel) or host (downer panel) cells were transgenic for *Cx3cr1gfp/+* (n=3-4 mice per group). Scale bar: 50μm **(J)** Experimental setup and confocal images depicting macrophages in regeneration ring of *Mrc1creERT2/creERT2 R26tdT/tdT*mice, induced with TAM in parallel to punch injury and analyzed 14 days later (n=3). Scale bar: 50μm. Data are mean±SEM. **p<0.01, ****p<0.0001 (one-way ANOVA, Tukey’s multiple comparison test if not stated otherwise). See also Figure S1.

### Rapid transcriptional adaptation of monocytes recruited to peripheral nerves

In order to distinguish recruited and resident sNAM after extraction from the tissue, we induced *Cx3cr1^creER/gfp^ R26^tdT/tdT^*mice before injury. Thereby, prior resident sNAM were labelled with both fluorescent reporter proteins, while recruited sNAM-like macrophages at injury site expressed only GFP (Fig. 1A, 2A). Interstitial macrophages do not express CX3CR1 under any of these conditions and thus were not labelled. We then sorted tdTomato^+^ GFP^+^ *bona fide* sNAM and double negative interstitial macrophages before and 10 days post injury. Only upon trauma, we additionally identified tdTomato^neg^ GFP^+^ cells, hence representing newly recruited sNAM-like macrophages (Fig. S2A). Using RNAseq, we found that the transcriptional differences of *bona fide* sNAM and interstitial macrophages persisted after injury, as samples extracted in homeostasis and after injury clustered closely together (Fig. 2B). sNAM exhibited a characteristic core transcriptional signature including genes such as *Axl*, *Bcl2a1* and *Cd9* as observed before (Kolter et al., 2019) (Fig. S2B). One of the highest upregulated genes in sNAM as compared to interstitial macrophages was *Ramp3*, the receptor of the neuropeptide calcitonin gene-related peptide (CGRP) (Fig. S2C). After injury, sNAM displayed 105 significantly upregulated genes, which were mainly associated with proliferation and DNA repair (Fig. 2D, S2D). Notably, 10 days post injury newly recruited sNAM-like macrophages displayed an intermediate transcriptional profile between sNAM and interstitial macrophages, in line with their transitional state in the tissue (Fig. 2B). Importantly, sNAM signature genes such as *Itgav*, *Bcl2a1* and *Cd72* were increased in sNAM-like macrophages as well (Fig. 2C). In contrast, genes that were significantly higher expressed in resident sNAM as compared to recruited sNAM-like cells, were associated with tissue residency e.g., the phosphatidylserine receptor gene *Timd4* (Fig. 2D). In comparison to interstitial macrophages, genes associated with cell migration and adhesion as well as neurogenesis were enriched (Fig. 2E). Signature genes such as *Axl, F11r* and *Ramp1/3* were similarly upregulated in resident sNAM and recruited sNAM-like macrophages after injury, whereas hallmark genes of interstitial macrophages such as *Mrc1* were consistently downregulated (Fig. 2F, G, S2E). Together, these data suggest that bone marrow-derived macrophages can converge with embryonic, strongly tissue-adapted macrophages *in vivo* with respect to immunophenotype and transcriptional profile upon opening of a novel, yet highly specific tissue niche.

**Figure 2.**
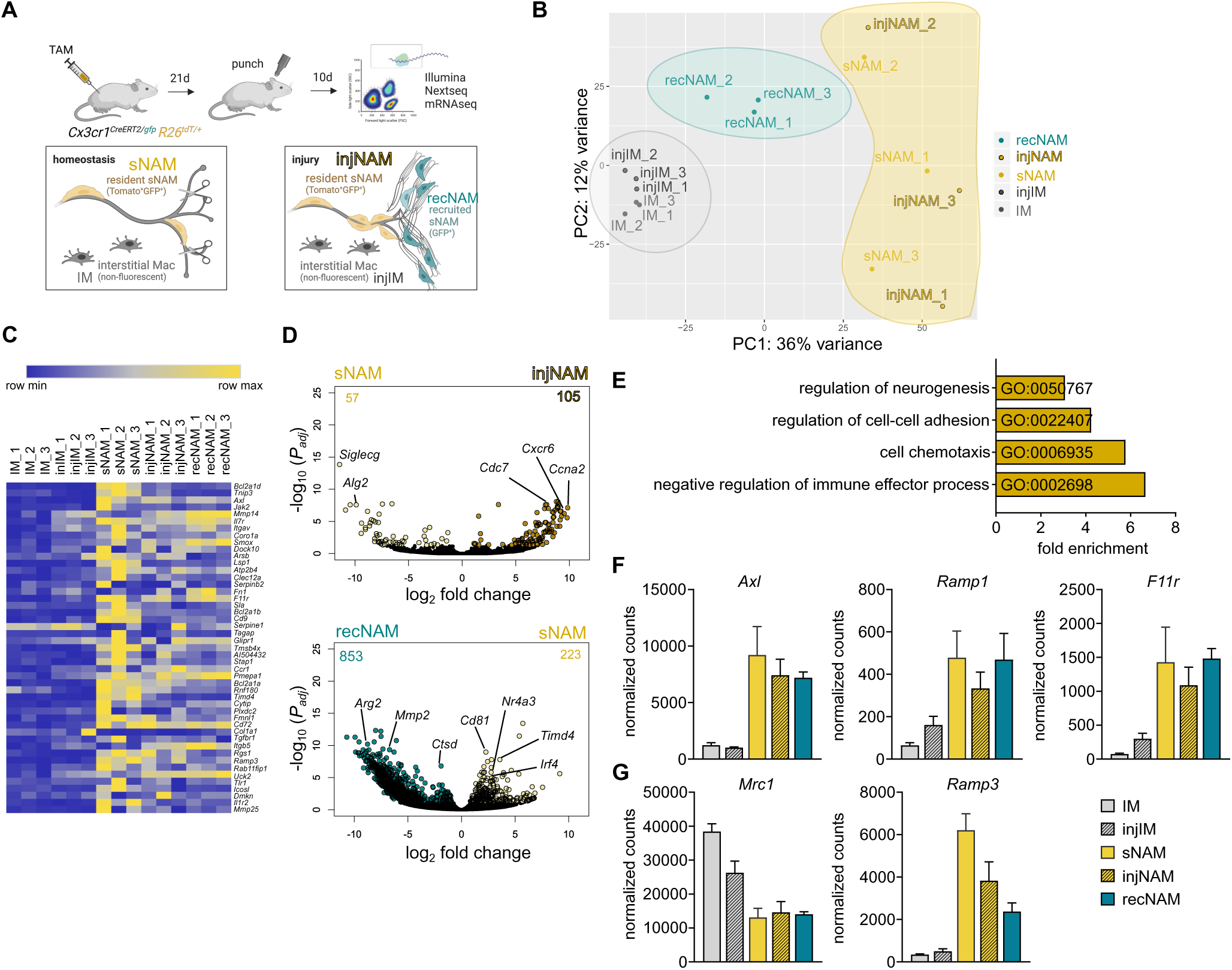
Rapid transcriptional adaptation of monocytes recruited to peripheral nerves. **(A)** Experimental setup to sort sNAM and interstitial macrophages (IM) from steady state and 10 days post injury (inj) for RNAseq. Mice were punched in the left ear (inj) 21 days after TAM induction, the left ear was left intact (steady state). Macrophages were sorted from both ears 10 days later and processed for RNAseq. RecNAM: Recruited sNAM. **(B)** Principal component analysis (PCA) of sorted macrophage subsets. **(C)** Heatmap depicting the top 50 upregulated “sNAM core” genes from Kolter et al, 2019. **(D)** Volcano plots of differentially expressed genes in sNAM upon injury. Yellow/green dots represent a log2-transformed fold change >1 and p≤ 0.05 (Padj = two-tailed Benjamini-Hochberg test P value). **(E)** GO biological pathways enriched in recNAM in comparison to sNAM (>1.5 fold change, FDR <0.01). **(F)** Normalized counts of selected genes similarly expressed in sNAM subsets. **(G)** Normalized counts of *Mrc1* and *Ramp3* in sorted macrophages subsets. Data are mean±SEM. See also Figure S2.

### Sensory neurons drive macrophage adaptation

In order to further dissect the interaction of macrophages and sensory neurons, we established an *ex vivo* system, where M-CSF-induced bone marrow-derived macrophages (BMDM) from *Cx3cr1^gfp/+^* mice were cocultured with sensory neurons differentiated from DRGs (Fig 3A). DRGs contain the cell bodies of dermal sensory neurons including diverse subtypes, which are characterized by expression of neuropeptides such as CGRP and substance P upon differentiation (Fig. S3A, B). We found that macrophages preferentially localized to neuronal somata and outgrowing neurites in the cultures, which was associated with cell morphological changes, i.e., an elongated cell body, and with physical alignment to axons (Fig. 3B). Moreover, live cell imaging revealed that macrophages showed patrolling behavior along the neurites (Fig. 3C, Movie S1) and exhibited strongly increased CX3CR1 expression, both of which was reminiscent of sNAM *in vivo* (Fig. 3B, D, E, S3C, D). Yet in contrast to sNAM in the tissue, CD206 was upregulated in macrophages in cocultures (Fig. 3D, E). F4/80 was significantly decreased, whereas the macrophage-defining surface markers CD11b, CD45 and CD64 were not affected (Fig 3E, S3E, F). Next, we sorted macrophages from cocultures and performed bulk RNAseq analysis. Macrophages cocultured with sensory neurons showed a substantially altered transcriptome with 5393 differentially expressed genes as compared to macrophages cultured alone (Fig. 3F, S3G). Gene ontology (GO) term enrichment analysis revealed upregulated gene sets associated with nerve development and differentiation (Fig 3G). Furthermore, macrophages from cocultures shared a common core signature with *bona fide* resident sNAM extracted from the skin, including 2169 differentially expressed genes such as *Axl*, *F11r* and *Gpr34* (Fig 3 F, H, I). Overall, these results demonstrate that contact with sensory neurons induces profound changes in phenotype and transcriptome of macrophages with substantial similarities *in* and *ex vivo*.

**Figure 3.**
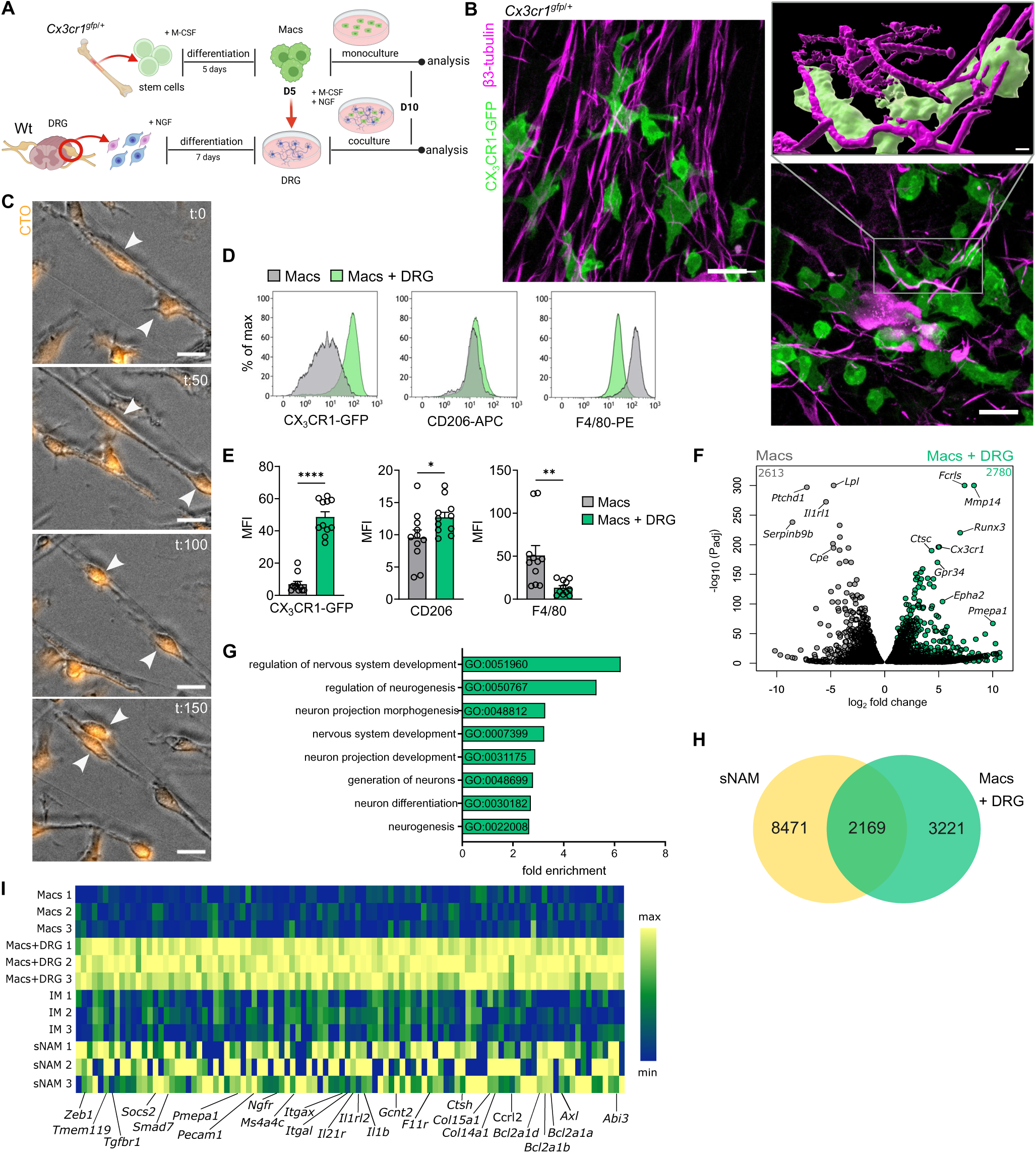
Sensory neurons drive macrophage adaptation. (A) Timeline of cocultures of Macrophages (Macs) and DRG cells. (B) Representative immunofluorescence (IF) staining of cocultures. Macs associated with axons (left) or neuron somata (bottom right). Zoom in: 3D reconstruction of Macs aligned along single neurites. Scale bars: 20 µm; zoom in: 3 µm. (C) Images from live cell imaging of cocultures after 3 days of culture, showing Macs moving along neurites (arrows). Macs were labelled with cell tracker orange (CTO) prior to coculturing. Time (t) is indicated in minutes. Scale bars: 20 µm. (D) Comparison of Mac surface marker after 5 days in mono- and coculture, measured by flow cytometry. (E) Analysis of surface marker expression measured by mean fluorescence intensity (MFI) of Macs in mono-and coculture (n= 11). (F) Volcano plot of differentially expressed genes in Macs in mono- and coculture. Grey/green dots represent log2-transformed fold change >1. Padj = two-tailed Benjamini-Hochberg test P value. (G) GO terms enriched in BMDM in coculture including “nerve” and “neuron”. (H) Venn diagram of DEG in Macs in coculture and sNAM *in vivo* compared to Macs in monoculture and interstitial macrophages in the dermis, respectively. (I) Heatmap of top 100 DEG upregulated in Macs in coculture with DRG *in vitro* compared to Macs in monoculture in RNA-seq. Expression of these genes was compared to sNAM and IM *in vivo* (Figure 2). Data are mean±SEM. *p<0.05, **p<0.01, ***p<0.001 (two-tailed unpaired t-test). See also Figure S3.

### TGF-β imprints nerve-associated macrophages

Next, we investigated signaling pathways activated in macrophages by the presence of DRG neurons *in vitro*. We found “response to TGF-β” and “TGF-β receptor signaling pathway” to be among the most prominently enriched GO terms (Fig. 4A, B). In particular, *Tgfbr1* and genes downstream of TGF-β signaling including *Smad3* and *Smad7* were upregulated along with genes under the regulatory influence of TGF-β such as *Gcnt2*, *Pmepa1* and *Runx3* (Fig 4C, D). Immunofluorescence staining and ELISAs revealed strong expression of TGF-β1 in macrophages and DRG cells, which was further increased in cocultures (Fig 4E, F). TGF-β is expressed in a wide range of tissues and has pleiotropic effects on different cell types. In macrophages, diverging effects of TGF-β have been reported, ranging from dampening of proinflammatory signaling to induction of migration (Batlle and Massague, 2019; Sanjabi et al., 2017). Moreover, TGF-β is well established to promote differentiation of alveolar macrophages (Yu et al., 2017) and embryonic microglia (Butovsky et al., 2014). However, the influence of TGF-β on skin macrophages remained largely unclear, in particular due to its context-dependent properties. Thus, we first treated BMDM with recombinant, activated TGF-β *in vitro* and found it to induce a strong concentration-dependent upregulation of CX3CR1 and concomitant downregulation of F4/80 and loss of CD206 (Fig 4G, S4A, B). Thus, TGF-β caused phenotypic changes in macrophages, which resembled those in cocultures with DRG neurons with the exception of CD206. Moreover, transcriptional profiling indicated enrichment of genes involved in cell adhesion and migration (Fig S3C). Notably, TGF-β-treated macrophages also exhibited transcriptional similarities with BMDM cocultured with DRG neurons, and upregulated sNAM signature genes, such as *Fcrls*, *F11r* and *Axl* (Fig 4H, S4D, E). We thus hypothesized that TGF-β imprinted macrophages in the neuronal environment. This rational was further supported by the finding that a selective inhibitor of the TGFBR1 kinase (SB-525334), which inhibits TGF-β-induced phosphorylation and nuclear translocation of SMAD2/3, prevented upregulation of CX3CR1 in macrophages cocultured with neurons from DRGs (Fig 4I, S4F). Furthermore, expression of the sNAM signature genes *F11r*, *Fcrls* and *Pmepa1* was suppressed in macrophages when TGF-β signaling was blocked (Fig S4G). Interestingly, the TGF-β signaling inhibition led to an even higher expression of CD206 in mono- and cocultures (Fig 4I, J, S4F, G). This observation indicated that additional factors in the cocultures led to initial upregulation of CD206, which was then further modulated by TGF-β. In agreement with this, macrophages treated with only TGF-β lost CD206 expression, reminiscent of sNAM *in vivo* (Fig. 4G, S4A). Finally, we genetically depleted *Tgfbr2* in macrophages by using bone marrow from *Mrc1^creERT2/creERT2^ Tgfbr2^flox/flox^* mice and OH-TAM treatment during differentiation of BMDM. In line with the pharmacological inhibition, we observed downregulation of CX3CR1 in macrophages in coculture with DRG neurons (Fig. S4H). Collectively, we identified TGF-β signaling as a key driver of immunophenotypic and transcriptional adaptation of macrophages in contact with peripheral nerves.

**Figure 4.**
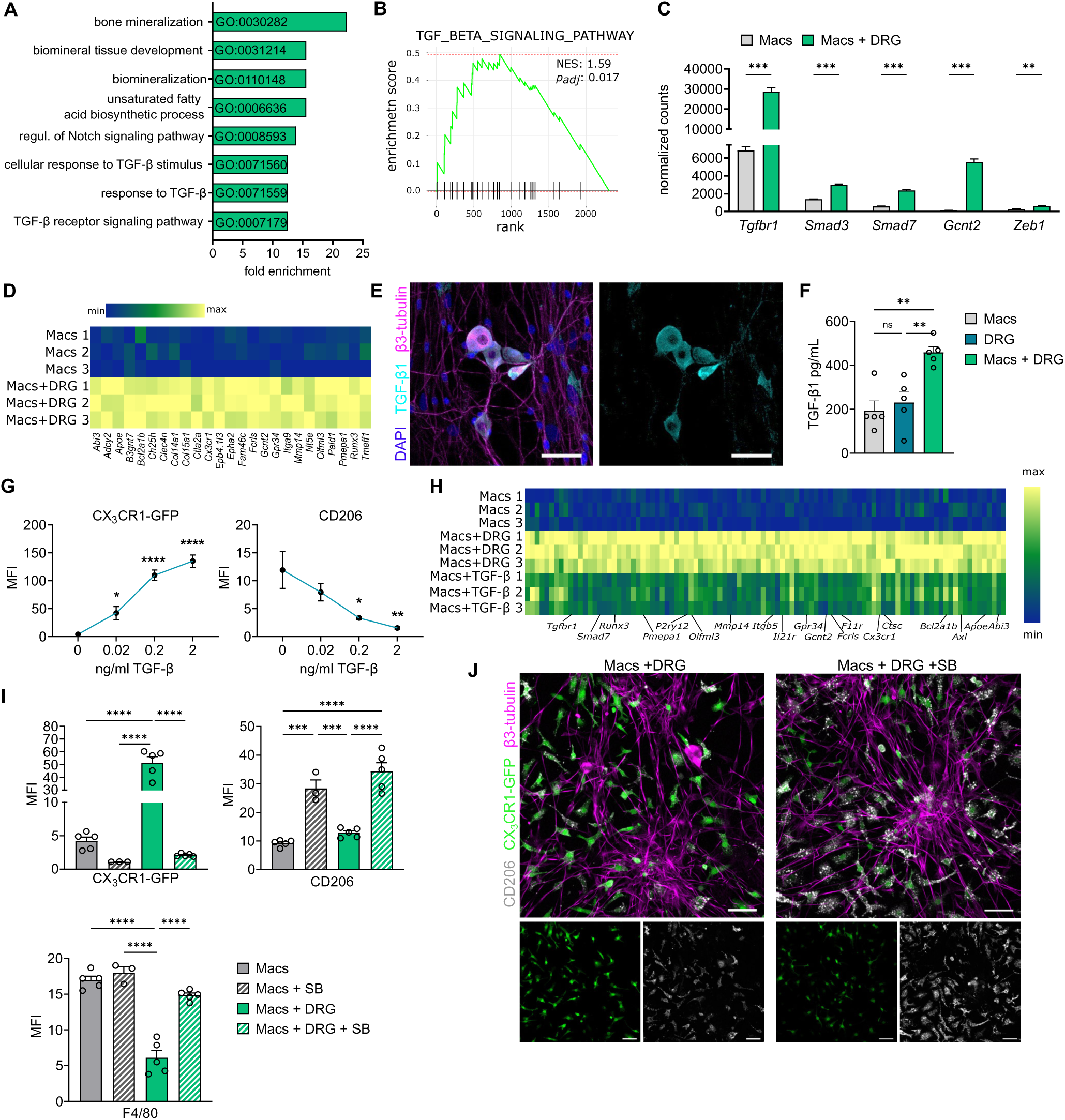
TGF-β imprints nerve-associated macrophages. (A) Gene ontology (GO) slim biological process pathways enriched in Macs cultured with neurons from DRG. Visualization of terms with highest fold enrichment in genes with a base mean >100. (B) Fast Gene Set Enrichment Analysis (fgsea) of ranked DEG (Mac in cocultures versus monocultures). Positive enrichment indicates genes of the “TGF_BETA_SIGNALING_PATHWAY”are enriched in Mac in cocultures. (C) Normalized counts of TGF-β related genes upregulated in RNAseq in cocultures (n=3). (D) Heatmap of top 25 genes related to TGF-β upregulated in co-cultured Macs in comparison to Macs cultured alone. (E) TGF-β1 staining of sensory neurons cultured from DRGs. Scale bar: 50 µm. (F) TGF-β1 ELISA of Macs and DRG in monocultures and coculture (n=5). (G) MFI of CX_3_CR1-GFP and CD206 in Macs cultivated for 5d with different concentrations of recombinant TGF-β, quantified by flow cytometry (n=3-5). Quantitative assessment of the influence of varied TGF-β concentrations on the expression of depicted surface markers in Macs after 5 days of treatment. (H) Heatmap of the 100 most enriched DEG upregulated in Macs treated with recombinant 0.02 ng/ml TGF-β and Macs in coculture with DRG (CC) compared to Macs in monoculture (Ctrl) after 5 days of culture. (I) MFI of Macs in mono-and cocultures treated with TGFBR1 inhibitor SB525334 (SB) or medium only for 5 days, measured by flow cytometry (n=3-5 per group). (J) Confocal microscopy of cocultures with or without SB525334 (SB) stained for CD206 and β3-tubulin. Scale bar: 50 µm. Data are mean±SEM. *p<0.05, **p<0.01, ***p<0.001, ****p<0.0001, ns, not significant (one-way ANOVA, Tukey’s multiple comparison test if not stated otherwise). See also Figure S4.

### TGF-β drives sNAM differentiation and self-maintenance in the skin

We next examined if sNAM exhibited a TGF-β-driven transcriptional signature *in situ*. Indeed, we found that sNAM showed high expression of *Tgfbr1* and the TGF-β target genes *Gcnt2* and *Zeb1*, both in steady state and after axonal injury (Fig. 5A). Moreover, genes driving the leading edge of the gene set enrichment analysis for TGF-β in macrophages cultured with neurons from DRG*in vitro* (Fig. 4B) (i.e., core members of the gene set with a potential biological relevance in macrophages) were also upregulated in CX3CR1^high^ sNAM in adult skin (Fig. 5B). Additionally, early in postnatal development (postnatal day 14), when most dermal macrophages still express CX3CR1 (Kolter et al., 2019), all dermal macrophages showed increased expression of TGF-β-related genes (Fig. 5B). This indicated that TGF-β globally steers early and potentially embryonic development of dermal macrophages, while the signature is only and highly selectively maintained in adult sNAM derived from prenatal sources, which may be due to microenvironmental cues. Since the cell bodies of sensory neurons innervating the skin are located in the DRGs, we then analyzed if DRG neurons produced TGF-β *in vivo*. In publicly available single-cell RNA-sequencing (scRNA-seq) datasets (Sharma et al., 2020), we indeed found expression of *Tgfb1* and *Tgfb2* in CGRP^+^ neurons and Aδ- and Aβ-low threshold mechanoreceptors (LTMR), respectively (Fig. 5C, S5A, B). In order to specifically – for the skin – target sNAM, we employed *Cx3cr1^creERT2/+^ R26^yfp/yfp^* (ctrl) mice. In line with the results of *Cx3cr1^creERT2/gfp^ R26^tdT/+^* mice (Fig. 1A), only skin macrophages associated with nerves (identified by co-localization with β3-tubulin, absence of CD206 expression and YFP expression) recombined after TAM 10 induction, in contrast to CD206^+^ interstitial macrophages and monocytes (Fig. S5C, S5D). Sorted YFP^+^ macrophages expressed sNAM signature genes such as *Fcrls*, *Axl*, *Plxna4* and *Ramp1-3* (Fig. S5E) and TGF-β related genes were strongly upregulated in comparison to YFP^-^ macrophages (Fig. S5F). Next, we crossed these mice with conditional *Tgfbr2^flox/flox^* mice in order to specifically deplete TGF-β signaling in sNAM. In notable support of TGF-β imprinting the sNAM phenotype, the conditional depletion of *Tgfbr*2 resulted in a loss of YFP^+^ macrophages associated with dermal nerves within 4 weeks (Fig. 5D, E, S5G). Remaining *Tgfbr2*-deficient YFP^+^ cells sorted from the skin showed reduced *Cx3cr1* and *Axl* expression (Fig. 5F). Moreover, *Tgfbr2* disruption strongly increased CD206 in the rather scarce YFP^+^ macrophages associated with sensory nerves in whole mount imaging (Fig. 5D, G). Collectively, these analyses suggest that sNAM throughout development rely on homeostatic TGF-β signaling both for self-maintenance and for expression of signature genes.

**Figure 5.**
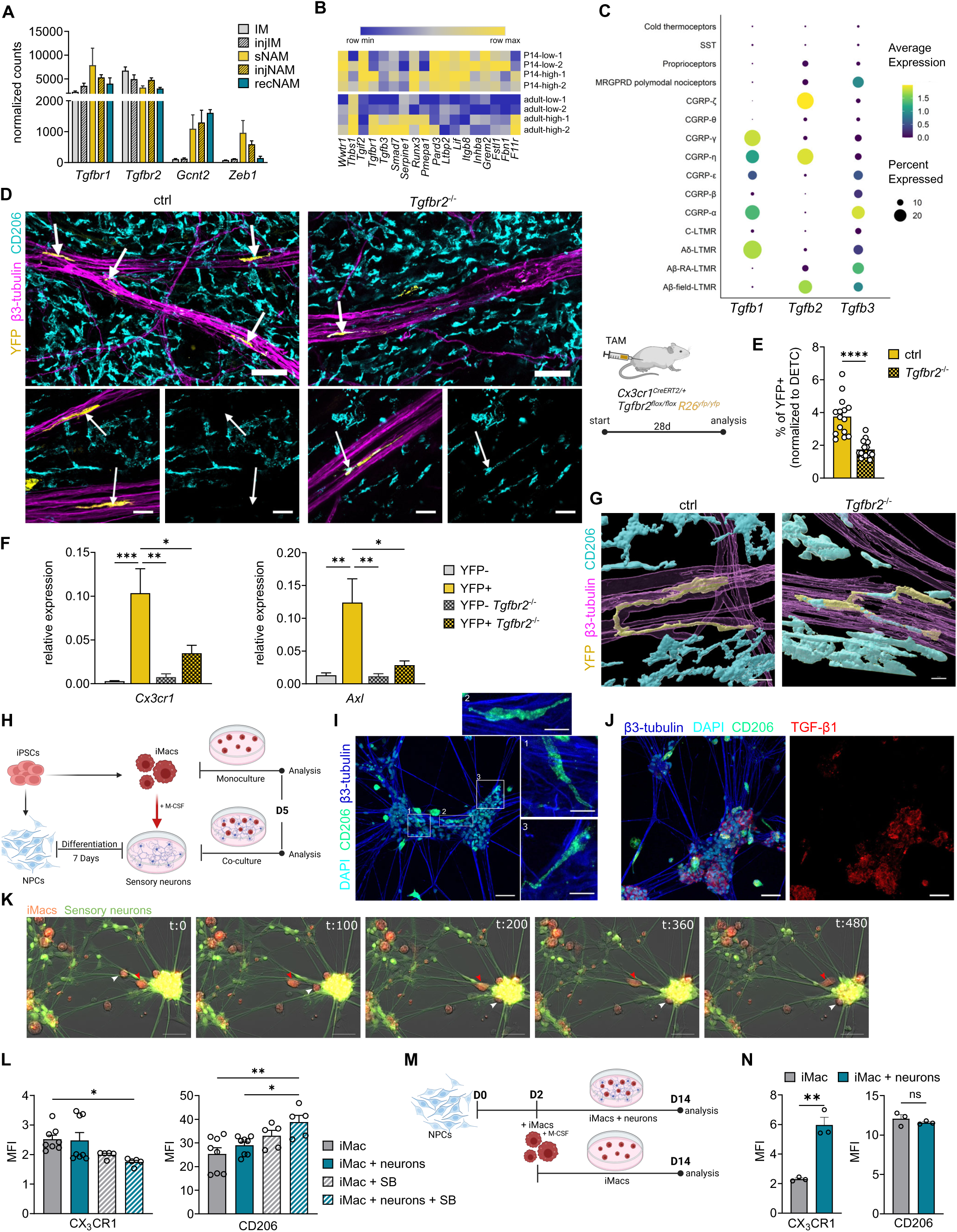
TGF-β drives sNAM differentiation and self-maintenance in the skin. (A) TGF-β-related genes in sorted dermal macrophages (normalized counts RNAseq). (B) Heatmap depicting genes, driving the leading edge of GSEA for TGF-β signaling in Fig. 4B, in CX_3_CR1^low^ and ^high^ macrophages in P14 and adult dermis (Kolter et al, 2019). (C) Dot plots of *Tgfb1-3* expression in neuronal subtypes of DRG. GSE19088 (Sharma et al, 2020 Nature), LTMR: low-threshold mechanoreceptor, CGRP: calcitonin-gene related peptide, SST (somatostatin+ pruriceptors). (D) *Cx3cr1^creERT2/+^ R26-yfp^flox/flox^Tgfbr2^flox/flox^*(*Tgfbr2^-/-^)* mice were induced with TAM 4 weeks prior to analysis. *Cx3cr1^creERT2/+^ R26-yfp^flox/flox^* (ctrl) were used as controls. Whole mount images depicting CD206 expression in remaining YFP^+^ nerve-associated macrophages in *Tgfbr2^-/-^* mice. Arrows point to nerve-associated macrophages. Scale bar: 50µm, zoom in: 20µm. (E) Quantification of YFP^+^ macrophages normalized to long-lived dendritic epidermal γδ T cells (DETC) in ear skin of mice described in (D) (n=15 per group). (F) Relative gene expression of depicted genes in sorted YFP^+^ and YFP^−^ macrophages sorted from mice in (D). n=6-7 per group. (G) 3D reconstruction of nerve-associated YFP^+^ macrophage in ctrl and *Tgfbr2^-/-^* mice (H) Timeline of human coculture with induced pluripotent stem cell (iPSC)-derived sensory neurons and iPSC-derived macrophages (iMacs). iPSC-derived sensory neurons were differentiated from neuron precursor cells (NPCs). (I) Confocal images of iMacs in coculture. White boxes (1-3): Elongated iMacs at ganglia-like structures of sensory neurons. Scale bar: 50 µm; zoom in: 20 µm. (J) Confocal image of ganglia-like structures in human coculture expressing TGF-β. Scale bar: 50 µm. (K) Images from live cell imaging of cocultures after 3 days of culture, showing iMacs moving along neurites (red arrowhead) and clustering at ganglia-like structures (white arrowhead). iMacs were labelled with cell tracker orange and sensory neurons with cell tracker green prior to co-culturing. Time (t) is indicated in minutes. Scale bars: 50 µm. (L) MFI of CX_3_CR1 and CD206 in iMacs in mono- and cocultures with and without TGFBR1 inhibitor SB525334 (SB) measured by flow cytometry (n=5-8). (M) Experimental setup. Human iMacs were added during differentiation (day 2) of iPSC-derived sensory neurons (n=3). (N) MFI of CX_3_CR1 and CD206 in iMacs cultured alone or together with iPSC-derived sensory neurons for 12 days, measured by flow cytometry (n=3) (two-tailed paired *t*-test). Data are mean±SEM. *p<0.05, **p<0.01, ***p<0.001, ****p<0.0001 (one-way ANOVA, Tukey’s multiple comparison test if not stated otherwise). See also Figure S5.

Next, we aimed to explore whether the specific role of TGF-β in creating a local macrophage niche was conserved from mice to humans. To assess this, we differentiated human skin fibroblast-derived induced pluripotent stem cells (iPSC) to sensory neurons, which formed dense neurite networks *in vitro* and had the capacity to secrete neuropeptides, such as CGRP and substance P (Fig. S5H) (Muller et al., 2018). iPSC-derived sensory neurons were then cocultured with human iPSC-derived macrophages (iMacs), which have been demonstrated to differentiate into tissue macrophages with genetically determined properties when exposed to organ-specific cues (Craig-Mueller et al., 2020; Takata et al., 2017) (Fig 5H). Similar to our observations in mouse cocultures, we found elongated iMacs in close proximity to sensory neurons, which patrolled along neurites and often accumulated at ganglia-like structures (Fig 5I, K, Movie S2). In cocultures with iMacs, these ganglia-like structures showed strong TGF-β expression, which was not observed when neurons were cultured alone (Fig. 5J, S5I). Indeed, various subtypes of nociceptors in human DRGs also expressed *Tgfb1* in a published spatial transcriptomic dataset (Fig. S5J) (Tavares-Ferreira et al., 2022). While CX3CR1 was only slightly elevated in cocultures with neurons, the inhibition of TGF-β signaling led to a decrease of CX3CR1 and an increase of CD206 in iPSC-derived human macrophages (Fig 5L, S5K, L). Finally, when iMacs were cocultured with NPCs already early during their differentiation into sensory neurons, they displayed strong upregulation of CX3CR1 in comparison to iMacs cultured alone (Fig. 5M, N, S5M). Collectively, these data suggest that the sNAM signature in human macrophages is also driven by TGF-β.

### Local activation of TGF-β by macrophages requires physical association with sensory neurons

TGF-β is involved in the development and/or survival of alveolar macrophages, epidermal Langerhans cells (LCs) and of embryonic microglia (Butovsky et al., 2014; Kaplan et al., 2007; Yu et al., 2017), yet it is thought to be dispensable for the self-maintenance of adult microglia and other tissue macrophages, including those in the dermis (Schridde et al., 2017; Yu et al., 2017; Zöller et al., 2018). We confirmed that microglia and colonic macrophages, which were targeted in our inducible system due to their high CX3CR1 expression, were indeed not quantitatively affected by abrogated TGF-β signaling (Fig. 6A). Additionally, and as opposed to sNAM, CX3CR1 and CD206 were expressed at normal levels (Fig. S6A, B). However, YFP^+^ microglia showed decreased *Fcrls* expression (Fig. S6B), in line with the published role of TGF-β in maintaining microglia homeostasis (Butovsky et al., 2014; Zöller et al., 2018). In contrast, alveolar macrophages and Langerhans cells were not targeted in this system in agreement with their lack of CX3CR1 and, accordingly, Cre expression (Yona et al., 2013). However, since CX3CR1 was one of the highest expressed genes in macrophages treated with TGF-β (Fig. 4G, H), we wondered why alveolar macrophages and LC, which are known to be exposed to TGF-β *in vivo*, generally do not express CX3CR1 in adult tissues. Notably, TGF-β has been shown to upregulate the master transcriptional regulator PPAR-γ in alveolar macrophages (Yu et al., 2017), yet we found PPAR-γ to be downregulated in BMDM cocultured with neurons or treated with activated TGF-β (Fig. S6C). In the lungs, GM-CSF produced by alveolar epithelial cells also induces PPAR-γ and is required for maintenance of alveolar macrophages (Schneider et al., 2014). In line with this notion, BMDM differentiated with GM-CSF instead of M-CSF and TGF-β completely lacked CX3CR1 expression (Fig. S6D). In addition, CX3CR1 expression was abrogated in fully differentiated BMDM when GM-CSF was added to the medium, even in the presence of TGF-β (Fig. 6B, S6E). Moreover, addition of GM-CSF to cocultures led to downregulation of CX3CR1 in macrophages cultured with DRG neurons while other markers were not affected (Fig. 6B, S6F). As we also did not observe downregulation of CX3CR1 nor upregulation of CD206 in microglia or colon macrophages upon *Tgfbr2* depletion (Fig. S6A, B), the immunophenotypic properties of TGF-β thus appear to be modulated by further niche-specific signals, such as other growth factors.

**Figure 6.**
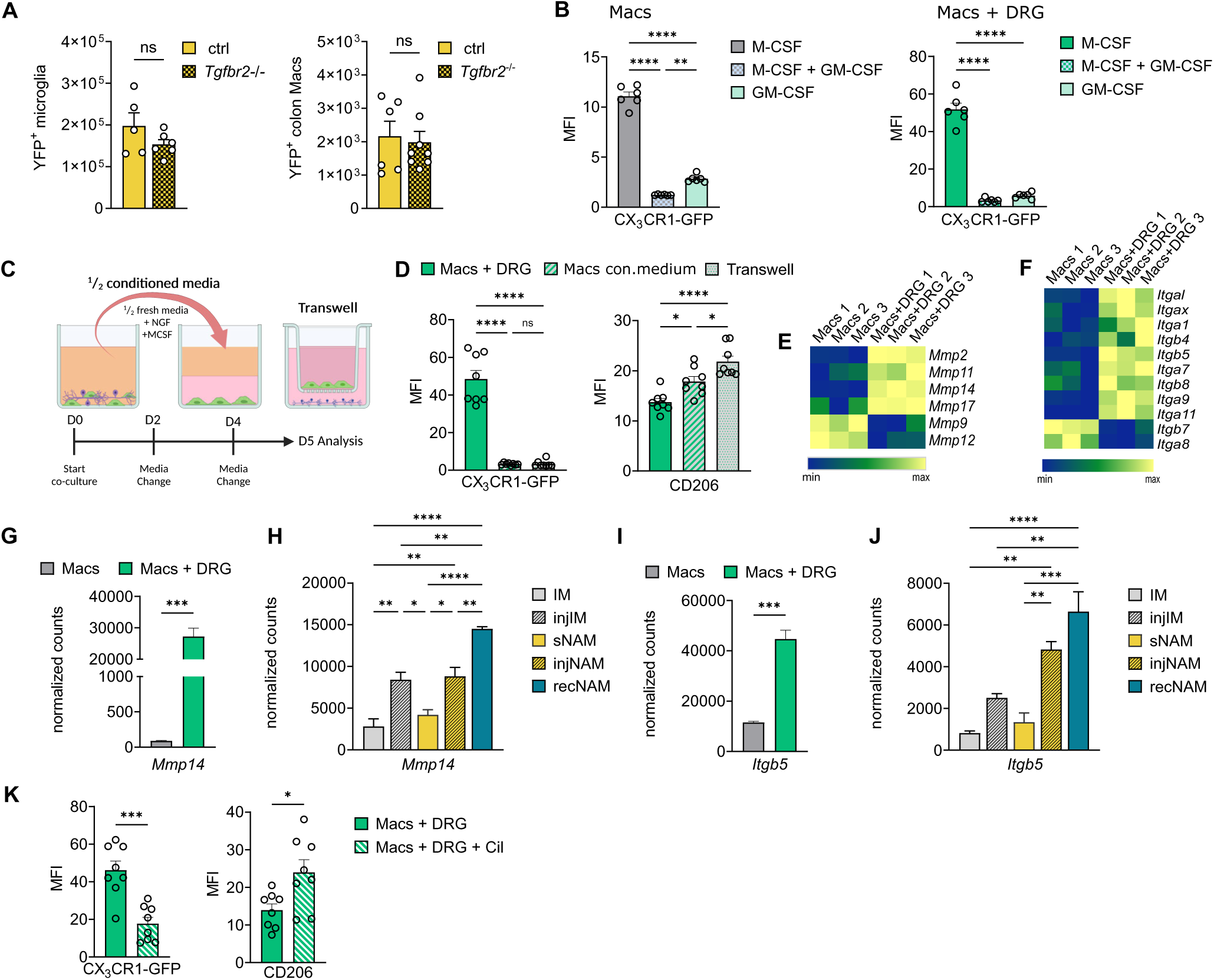
Local activation of TGF-β by macrophages requires physical association with sensory neurons. (A) Absolute counts of YFP^+^ microglia (CD45^lo^ CD11b^hi^ CD64^hi^ in brain and of colonic macrophages (CD45^hi^ CD11b^hi^ CD64^hi^ Lin^neg^) in mice 4 weeks post TAM (two-tailed unpaired *t*-test). (B) Comparison of CX_3_CR1 expression of Macs in mono- and cocultures in the presence of either M-CSF, GM-CSF or both, measured by flow cytometry (n=6). (C) Experimental set up for conditioned (con.) medium and transwell plate approaches. (D) MFI for CX_3_CR1 and CD206 in Macs either cocultured with DRGs, treated with conditioned medium (con. medium) from cocultures or cultured in transwells with DRG (n=8). (E) Heatmap of DEGs for metalloproteases in Macs in mono-and cocultures. (F) Heatmap of DEGs for integrins in Macs in mono-and cocultures. (G) Normalized counts of *Mmp14* in Macs in mono- or cocultures (two-tailed unpaired *t*-test). (H) Normalized counts of *Mmp14* in dermal macrophages of RNAseq data from Figure 2. (I) Normalized counts of *Itgb5* in mono- or cocultures (two-tailed unpaired *t*-test). (J) Normalized counts of *Itgb5* in dermal macrophages of RNAseq data from Figure 2. (K) Quantification of Mac surface marker expression in cocultures treated with ανβ5 inhibitor cilengitide (Cil) (two-tailed unpaired *t*-test) (n=8).

In extension of this concept, it seems notable that sNAM represent a minor population (2-3%) in the dermis with a highly specific TGF-β-dependent phenotype. In contrast to sNAM, the majority of dermal macrophages lacks CX3CR1, but instead shows strong expression of CD206. This situation is highly dissimilar to that of alveolar macrophages and LC, both of which predominate in their respective tissues and show rather homogenous immunophenotypes. This implies that TGF-β must act with highest precision and very locally in the neuronal niche in order to mediate discrepant effects on sNAM and immediately adjacent interstitial macrophages. Accordingly, we addressed whether macrophages require physical contact to neurons by culturing BMDM either with conditioned medium obtained from cocultures or in a transwell system, where a membrane with 0.4 μm pores prevented physical contact of BMDM and sensory neurons, but allowed the passage of soluble factors (Fig 6C). Interestingly, we found direct contact to be required for the induction of the sNAM-like phenotype in macrophages (Fig 6D, S6G). TGF-β is secreted as a biologically inactive complex with the disulfide-linked latency-associated peptide (LAP), which needs to be cleaved in order to enable binding to the TGF-β receptor. Latent TGF-β can be activated by enzymatic cleavage, e.g., via certain matrix metalloproteases, or by mechanical force upon binding of integrins to the LAP domain (Batlle and Massague, 2019). Consistent with this mechanism, in BMDM cocultured with neurons, we observed upregulation of genes typically involved in the interaction of macrophages with the extracellular matrix such as genes for metalloproteases and integrins (Fig 6E, F). Notably, the gene for the transmembrane metalloprotease *(Mmp14*), which is known to activate TGF-β, was upregulated both *in vitro*, i.e., in BMDM cocultured with neurons as well as in sNAM after injury *in vivo* (Fig 6E, G, H). In addition, the gene encoding the subunit of integrin subunit β5 (*Itgb5*), was highly expressed under these conditions (Fig 6F, I, J). Integrin subunit β5 builds a cell surface heterodimer with the integrin subunit αv. Treatment of the cocultures with a peptide inhibitor of integrin αvβ5 (Cilengitide) strongly decreased CX3CR1 and simultaneously increased CD206 expression (Fig 6K, S6H), indicating that integrins mediated local TGF-β activation when macrophages were physically associated with neurons. Thus, TGF-β acted in a spatially restricted fashion on macrophages, thereby modulating macrophages directly recruited to neurons, while sparing more distant cells. Moreover, the concerted action with tissue-specific factors such as GM-CSF induced a context-specific response of macrophages in different tissues.

### TGF**-**β signaling in macrophages impacts on neuronal growth

Macrophages support axonal outgrowth and repair after injury (Cattin et al., 2015; Kolter et al., 2019). Accordingly, we next assessed the effect of TGF-β signaling in sNAM on the resprouting of nerves after injury and induced *Cx3cr1^creERT2/+^ R26^yfp/yfp^ Tgfbr2^flox/flox^* (*Tgfbr2^-/-^*) and the same mice without the floxed gene (ctrl) with TAM and performed ear punches (Fig. 7A). Under these conditions, resident sNAM proliferate on injured axons immediately adjacent to the injury site and phagocytose myelin (Kolter et al., 2019). In *Cx3cr1^creERT2/+^ Tgfbr2^flox/flox^* mice, we found YFP^+^ nerve-associated macrophages to be reduced, indicating that TGF-β signaling was required for their self-maintenance and accumulation under injury (Fig. 7B, S7A). Furthermore, remaining YFP^+^ macrophages at axonal injury sites expressed high levels of CD206 which was not observed in control mice (Fig. 7C). However, the resprouting of injured axons was not affected (data not shown). We hypothesized that recruited sNAM functionally compensated for the loss of prior resident sNAM, since newly recruited sNAM, which upregulate CX3CR1 upon association with nerves *after* injury, were not targeted by this approach. Thus, we improved targeting of these sNAM-like macrophages by continuous administration of TAM (Fig. 7D). Thereby, both prior resident and macrophages newly recruited into the sensory nerve niche after the injury displayed high recombination rates. Notably, interstitial dermal macrophages and monocytes did not express YFP under continuous TAM treatment (data not shown, Fig. S7B). Less Ly6C^hi^ macrophages and neutrophils were observed at the site of injury in *Cx3cr1^creERT2/+^ Tgfbr2^flox/flox^* mice, although neutrophils and monocytes were slightly elevated in peripheral blood of the latter (Fig. 7E, F, S7C, D). In control mice, regeneration rings of resprouted nerves were populated by YFP^+^ macrophages (Fig. 7G). Most notably, recruited YFP^+^ macrophages at injury site did not associate with nerves and did not acquire the typical sNAM morphology in *Cx3cr1^creERT2/+^ Tgfbr2^flox/flox^* mice (Fig. 7G, H, S7E). Moreover, nerve resprouting was significantly impaired when TGF-β receptor signaling was lost in CX3CR1^+^ macrophages (Fig. 7G-I). Collectively, these data suggest that the local sensing of TGF-β in macrophages is required for their precise and spatially confined adaptation to the sensory nerve niche, and that this functional commitment of highly specialized macrophages is a prerequisite for tissue reinnervation upon injury.

**Figure 7.**
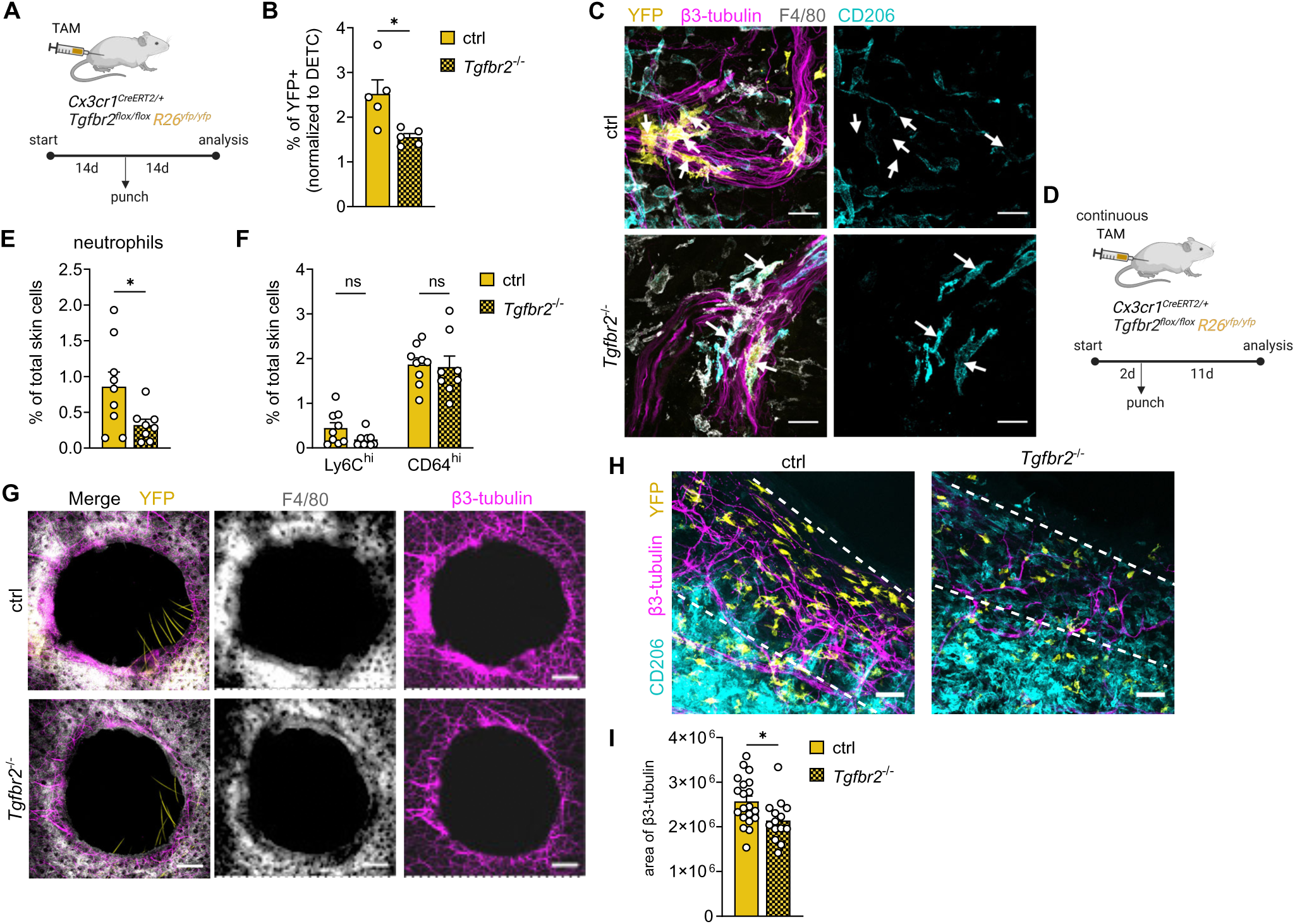
TGF-β signaling in macrophages impacts on neuronal growth. **(A)** Experimental setup to deplete *Tgfbr2* in sNAM in punch injury. **(B)** Quantification of YFP^+^ macrophages normalized to DETC in ear skin of mice. n=5 per group in 3 independent experiments. **(C)** Representative whole mount image depicting YFP+ macrophages recruited to injured nerve endings in mice described in (A). Scale bar: 50µm **(D)** Experimental setup to deplete *Tgfbr2* both in resident and recruited sNAM in punch injury. n=8-9 per group in 3 independent experiments. **(E)** Quantification of Ly6G^+^ neutrophils in the skin via flow cytometry. **(F)** Quantification of Ly6C^hi^ and CD64^hi^ cells in the skin 11 days post injury. **(G)** Whole mount images of resprouting nerves and associated sNAM at injury sites 11 days post injury. **(H)** Whole mount images depicting YFP^+^ macrophages in regeneration ring in control and *Tgfbr2^-/-^* mice. **(I)** Quantification of β3-tubulin^+^ area at injury sites of mice in (D). Data are mean±SEM. *p<0.05 (two-tailed unpaired *t*-test). See also Figure S7.

## DISCUSSION

Resident immune cells contribute to tissue homeostasis by sensing danger signals and mediating their timely clearance in order to avoid tissue destruction. Peripheral nerves transmit sensory information from tissues and also regulate inflammatory responses, which in turn affect neural signaling. The immune system and the nervous system thus need to interact and communicate to synchronize these processes and maintain tissue integrity as well as to avoid potential pathogenic dysregulation. Here, we uncover that macrophages are recruited to sensory nerves upon niche availability, which emerges for example after nerve injury. The physical interaction with sensory nerves drives a transcriptional program and long-lasting differentiation to sNAM in the tissue, which involves TGF-β as a molecular link propagating bi-cellular communication, ultimately enabling nerve regeneration.

Neuro-immune interactions have received increasing attention in recent years (Enamorado et al., 2023; Hoeffel et al., 2021; Pinho-Ribeiro et al., 2023; Zhang et al., 2021). In various tissues, macrophages have been found to associate with peripheral nerves, e.g., the enteric nervous system or sympathetic nerves in the gut or adipose tissue (De Schepper et al., 2018; Pirzgalska et al., 2017; Wolf et al., 2017). The phenotype and function of these macrophages strongly depends on the tissue and the type of nerve, e.g., macrophages in the enteric nervous system express β2-adrenergic receptors and mediate neuronal protection upon enteric infection (Gabanyi et al., 2016; Matheis et al., 2020). Macrophages thus have a remarkable capacity to adapt to their immediate environment and contribute to its proper function. The molecular cues governing this specification remain to be established. By combining selective gene targeting in the skin with cocultures, we essentially expand existing models on how macrophages are guided in neuronal niches: TGF-β signaling in combination with physically direct cell-cell-contact paves intratissular differentiation of macrophages with a very distinct phenotype and function which is required to support neuronal regeneration. This finding is in accordance with very recent observations by Viola et al. who found TGF-β to imprint muscularis macrophages in the intestine in the proximity of nerves and loss of TGF-β signaling in muscularis macrophages to reduce neuronal density in the intestine (Viola et al., 2023). However, nerve-associated macrophages in the intestine could not be specifically targeted in this study, due to the lack of a specific marker of these cells. The combination of our finding with that of Viola et al. indicates that TGF-β signaling may be a conserved factor supporting macrophages in contact with peripheral nerves, which then patrol these important intratissular structures with high fidelity. TGF-β is interesting in this respect, as it is secreted in a latent pro-TGF-β form and requires local activation. The latent complex is typically bound to cell surfaces via so-called milieu molecules, such the glycoprotein A repetitions predominant protein (GARP/LRRC32) or LRRC33, or to the extracellular matrix via latent TGF-β binding protein (LTBP) (Batlle and Massague, 2019; Qin et al., 2018). In particular, interaction with cognate integrins in combination with cytoskeleton-associated traction force enables a localized release. Thus, TGF-β action can be spatially extremely confined, relying on activation by highly regulated integrin-expression on specific cell subsets, and not on the rather ubiquitous presence of diffusible TGF-β. A similar mechanism has previously been shown for the CNS, where integrin-bearing astrocytes and microglia expressing milieu molecules cooperatively activate TGF-β with little spread of TGF-β activity to neighboring microglial cells (Qin et al., 2018). This model provides a satisfying mechanistic explanation on how macrophages are imprinted in a highly local fashion within the dermis, where sNAM and phenotypically highly distinct interstitial and perivascular macrophages coexists site by site. While we found that integrin β5 contributes to TGF-β activation in vitro, it is also conceivable that autocrine TGF-β, which is produced by all dermal macrophages (data not shown) is locally cleaved by proteases from sensory neurons or Schwann cells. In the CNS, it was recently suggested that integrin β8 expressed by radial glia may activate autocrine TGF-β1 from microglia during development (McKinsey et al., 2023). Potential additive effects in the PNS by autocrine TGF-β1 thus remain to be established in detail.

In contrast to fully differentiated sNAM, adult microglia do not depend on TGF-β for their survival and self-maintenance (Buttgereit et al., 2016; Zöller et al., 2018). Yet, TGF-β is required to maintain the homoeostatic microglia signature in adult mice and for their development in the embryo (Butovsky et al., 2014; Buttgereit et al., 2016; Zöller et al., 2018). On the other hand, alveolar macrophages critically depend on autocrine TGF-β for differentiation and survival via PPAR-γ (Yu et al., 2017). Notably, TGF-β was considered to be unique in its role for alveolar macrophages and Langerhans cells, but not for other tissue macrophages including dermal macrophages (Lambrecht, 2017; Yu et al., 2017). In contrast, by using more specific tools like inducible *in vivo* signal disruption, we have uncovered that TGF-β is also relevant for the local imprinting and differentiation of a highly dedicated dermal macrophage subset. Tissue heterogeneity thus needs to be accounted for when tissue macrophages are studied in depth. As peripheral nerves, similar to the CNS, require protection from external stressors and inflammation, TGF-β may here act as an important local immunosuppressor, regulating the phenotype and immune response of incoming and resident macrophages. In combination with further niche-specific cues, TGF-β can then shape the local identity of specific macrophages subsets leading to context-dependent transcriptional programs. Underlying molecular mechanisms may be conserved across sites and modified for instance by growth factors, which promote macrophage maintenance in the respective niche and orchestrate differential gene networks in combination with TGF-β. As an example, TGF-β leads - in concert with GM-CSF - to downregulation of CX3CR1 in macrophages.

Importantly, we introduce the coculture system with either mouse or human cells as a model to study neuro-immune interactions, phenocopying major characteristics of the interaction of sensory nerves and macrophages in the tissue. As the quantity of sNAM is very low and typically represents a few hundred cells per individual, this model enables valuable insights into sNAM behavior and underlying molecular mechanisms, including the opportunity to interfere with putative signaling pathways in a cell type-specific fashion. Thus, we complement very recently developed models, e.g., on the interaction of nociceptors with dendritic cells (Hanc et al., 2023). The ear punch injury model enables the investigation of macrophage adaptation to axons in case of niche opening in a kinetic fashion *in vivo*. We hypothesize that mechanisms uncovered in this model may be involved in the development of skin innervation during fetal development. As skin innervation starts at E13.5, interference with molecular pathways or cellular exchange would have to be performed during pregnancy, a challenging, albeit highly promising outlook for future studies. Moreover, it remains to be established how peripheral glial cells such as Schwann cells in the skin or satellite glial cells in the DRG contribute to the microenvironmental imprinting of macrophages. Finally, the function of CX3CR1 itself in sNAM remains to be solved. So far we did not observe an effect of CX3CR1 on proliferation or survival (data not shown), which has been demonstrated in arterial macrophages or DRG macrophages in peripheral nerve injury (Ensan et al., 2015; Guimaraes et al., 2023). Interestingly, TGF-β in skin wounds induces dermal dendritic cells to produce IL-31 which in turn activates sensory neurons and induces wound itching (Xu et al., 2020). Investigating the role of resident macrophages including their TGF-β signaling in skin disorders and neuroinflammatory conditions will thus be essential.

The peripheral nervous system is a vital component of healthy tissues and has, in contrast to the CNS, the ability to regenerate. Upon injury, the proximal ends of lesioned axons form growth cones with a limited potential to resprout. The cues driving regeneration are incompletely understood, however, peripheral immune cells are likely to contribute (Wofford et al, 2022). In the ear punch model, tissue-resident T cells were shown to positively influence nerve regeneration upon colonization by *Staphylococcus aureus* (Enamorado et al, 2022). Thus, it is tempting to speculate that macrophages in close contact with sensory nerves and primary bacterial sensors might contribute to this effect as well.

In summary, we found that the functional tissue macrophage phenotype was imprinted by the direct microenvironment. Macrophage survival and maintenance depended on specific locally supplied factors and on the availability of the cellular context of the respective tissue niche. Originally monocyte-derived macrophages readily integrated in the nervous niche when new axons sprouted, suggesting that – under homeostatic conditions - incoming monocytes were rather limited by the existence of self-maintaining resident macrophages than a principal lack of plasticity. In addition, we identified non-redundant function of sNAM for cutaneous nerve repair. Whether subtle differences between prenatally seeded sNAM and recruited sNAM-like cells originating from hematopoietic stem cells persist in the long-term remains to be established. Overall, a better understanding of the complex immune-mediated and thus potentially druggable functions of macrophages in the regeneration of the peripheral nervous system holds promise for the therapy of peripheral nerve injuries and neuropathies.

## ACKNOWLEDGEMENTS

We are indebted to Anita Imm and the Lighthouse facility of the University Medical Center Freiburg for their assistance with cell sorting and confocal microscopy, and to Marco Prinz (Freiburg) and Steffen Jung (Israel) for provision of mice. We thank the Center for Experimental Models and Transgenic Service (CEMT) of the University Medical Center Freiburg for excellent technical support. J.K. was funded by the Hans A. Krebs Medical Scientist Programme (Faculty of Medicine, University of Freiburg) and by the German Research Foundation (DFG) within the TRR359 (Project ID 491676693). P.H. received funding by the DFG (HE3127/9, HE3127/12, HE3127/16, TRR359 (Project ID 491676693), TRR167 (Project ID 259373024) and CRC1160 (Project ID 256073931). V.F. is employed by the Centre National de la Recherche Scientifique and is the recipient of grants from the Agence Nationale de la Recherche (ANR): ANR/ERAPerMed BATMAN (ANR-18-PERM-0001), ANR AUTOMATE (ANR-20-CE15-0018-01), and ANR LabCom INCREASE (ANR-22-LCV2-0009-01). S.B. was funded by the Hans A. Krebs Medical Scientist Programme (Faculty of Medicine, University of Freiburg). L.D. was supported by ANR/ERAPerMed BATMAN. B.V. was supported by a Marie Slodowska-Curie Individual Fellowship (H2020-MSCA-IF-2019 896095 VirIVITES). N.L. received funding from the European Research Council (ERC) under the European Union’s Horizon 2020 research and innovation program (grant agreement no. 852178) and is also funded by the European Union (grant agreement no. 101100859). Views and opinions expressed are, however, those of the author(s) only and do not necessarily reflect those of the EU or the ERC. Neither the EU nor the granting authority can be held responsible for them. N.L. was further funded by the DFG under Germany’s Excellence Strategy - EXC 2155 - project number 390874280. KK was supported by the DFG by project grants within the TRR359 (Project ID 491676693), TRR167 (Project ID 259373024), CRC1479 (Project ID 441891347), CRC1160 (Project ID 256073931) and by the DFG under Germany’s Excellence Strategy (grant no. CIBSS—EXC-2189, Project ID 390939984). Experimental schemes were created with Biorender.com.

## AUTHOR CONTRIBUTIONS

J.K. and C.D. conceptualized the study, designed experiments, analyzed the data and wrote the manuscript. J.K., C.D., S.B., R.A., Z.M.M., V.G. and F.L. performed experiments, G.V.L.S. and C.E.A.S. analyzed published scRNA-seq datasets, T.B. and N.L. generated iPSC-derived macrophages, B.V., L.D. and V. F. generated iPSC-derived neuron precursor cells. K.K. provided mice and critical revision. J.K. and P.H. supervised the project and acquired funding.

## METHODS

### CONTACT FOR REAGENT AND RESOURCE SHARING

Further information and requests for resources and reagents should be directed to and will be fulfilled by the Lead contact, Julia Kolter.

## DATA AND CODE AVAILABILITY

RNAseq data will be published on a database repository before publication.

## EXPERIMENTAL MODEL AND SUBJECT DETAILS

### Mice

All mice were on C57BL/6 genetic background. C57BL/6J and C57BL/6N mice were purchased from Jackson Laboratories (USA) or Charles River Laboratories (Germany). *Cx3cr1^gfp/+^* mice were a kind gift of Steffen Jung (Weizmann Institute, Rehovot, Israel). *Cx3cr1^creERT2^* mice were described in (Yona et al., 2013). For fate mapping analysis, mice were crossed to either floxed *Rosa26-*YFP (*R26^yfp/yfp^*) or *Rosa26-tdTomato* (*R26t^tdT/tdT^*) mice and *Cx3cr1^gfp/+^* mice in second generation to obtain of *Cx3cr1^creERT2/gfp^ R26-tomato* mice. *Mrc1^creERT2/creERT2^ R26t^tdT/tdT^* and *Cxcr4^creERT2/+^ R26t^tdT/tdT^* mice were obtained from Marco Prinz (University of Freiburg, Germany). *Mrc1^creERT2/creERT2^*and *Cx3cr1^creERT2^ R26^yfp/yfp^* mice were bred to *Tgfbr2^flox/flox^* mice provided by Katrin Kierdorf (University of Freiburg, Germany). *Cd45.1* and *beta-actin-EGFP* mice were purchased from Jackson Laboratories (USA). Mice were bred under specific pathogen-free conditions in the animal facilities of the University of Freiburg and housed in groups of up to five mice. Mice were kept in 12h light/dark cycles and food and water was provided *ad libitum*. Adult mice were between 6 and 20 wk of age if not mentioned otherwise. Both male and female mice were used for experiments, for transplantations mice were sex-matched. Littermates were randomly assigned to experimental groups. All animal experiments were approved by the Federal Ministry for Nature, Environment and Consumer’s protection of the state of Baden-Wuerttemberg.

## METHOD DETAILS

### Animal models and *in vivo* interventions

Cre recombinase was induced as previously described (Kolter et al, 2019). Briefly, 4 mg TAM were dissolved in 200 µl corn oil (both Sigma) and injected subcutaneously at 4 injections sites in mice under isoflurane narcosis. The treatment was repeated after 48 h. For bone marrow transplantations, mice were anesthetized with Ketamine and Xylazin. One ear of each recipient mouse was shielded with a customized lead device and lethally irradiated with 9 Gy. Subsequently, mice received 1×10^7^ donor-derived bone marrow cells i.v.. For ear punches, mice were punched in the center of the ear pinnae using a sterile stainless steel 2-mm hole punch (Type 292-2, Zoonlab, Germany).

### Tissue preparation

Dermal macrophages were isolated as described recently (Forde and Kolter, 2024). Briefly, mouse ears (dermis and epidermis) were minced into small pieces and subjected to enzymatic digestion by Dispase (0.25 U/ml; STEMCELL Technologies), Collagenase II (1 mg/ml; Worthington), and DNase I (0.04 mg/ml; Roche) in HBSS with 5% FCS at 1400rpm at 37°C for 2h. For colon macrophages, the colon was opened longitudinally and rinsed. Epithelial cells were dissociated by shaking twice for 15 min at 37°C in 2 mM EDTA and 10 mM HEPES in HBSS. Remaining tissue was washed, minced, and digested three times for 15 min at 37°C with 0.3 mg/ml Collagenase IV (Worthington), 5 U/ml Dispase (Corning), and 0.5 mg/ml DNase I (Roche) in HBSS supplemented with 2 % FCS. After digestion, the samples were filtered with a 70µm cell strainer.

Peripheral blood was withdrawn from the retro-orbital sinus and red blood cells were lysed with RBC lysis buffer (eBioscience). For microglia isolation, mice were transcardially perfused with ice-cold PBS. Brains were removed, homogenized using a Dounce homogenizer (Carl Roth) and the cell suspension was filtered through a 100 µm cell strainer. Myelin was removed by resuspension in 37 % Percoll (Sigma Aldrich) and centrifugation for 20 min at 300*g* (RT) without acceleration and break. To harvest neurons, DRGs from all spine sections were collected. In the first step of digestion, DRGs were incubated for 60 min in 10 mg/ml collagenase II dissolved in sterile Hanks Balanced Salt Solution (HBSS). In the second step, DRGs were digested with 0.25 % Trypsin (Gibco) for 20 min. Subsequently, DRGs were triturated by repeated pipetting in DMEM F12 (Gibco) medium and filtered through a 70 µm strainer and centrifuged for 5 min at 200 g.

### Cell cultures

For BMDM culture, bone marrow from femur and tibia was extracted and plated in DMEM medium, containing 10 % fetal bovine serum (FBS), 20 ng/mL M-CSF (Peprotech) and 0.05 % Ciprofloxacin (Frisenius Kabi). After five days of differentiation, BMDM were harvested. For DRG cultures, pelleted DRG cells were seeded in DMEM F12 and Neurobasal (Gibco) medium (1:1) containing 10% horse serum (Gibco), 10 ng/ml human NGF (Peprotech) and 0.05 % Ciprofloxacin in poly-D-lysin (Sigma Aldrich)-coated wells. Medium was changed every third day. Cells were treated for 10 min at 37°C with accutase solution (Sigma Aldrich) for harvesting.

### iPSC-derived cells

#### Human iPSC-derived macrophages

Human iPSC-derived macrophages were generated as described previously (Ackermann et al., 2022). iPSC cells were cultured on Geltrex^TM^-coated tissue culture flasks using E8 medium (containing DMEM/F-12 (GIBCO, Life Technologies), 64 mg/L ascorbic acid 2-phosphate, 14 µg/L sodium selenite, 543 mg/L, NaHCO3, 20 mg/L insulin and 10.7 mg/L human recombinant transferrin (all from Sigma-Aldrich) supplemented with 100 ng/mL hbFGF and 2 ng/mL hTGFß (both from Peprotech) under standard humidified conditions at 37 °C and 5 % CO2. The cells were split twice a week using Accutase^TM^ (Stemcell Technologies) and under the addition of 10 µM Y-27632 (Tocris, Bristol, UK). A complete medium change using Y-27632-free medium was performed 48 h after splitting. In order to start mesoderm priming, 5 x 10^5^ iPS cells were seeded in 3 mL of mesoderm priming I medium (E8 medium supplemented with 10 µM Y-27632, 50 ng/mL hVEGF and hBMP4 and 20 ng/mL hSCF (all from Peprotech) using CELLSTAR 6-well plates placed on an orbital shaker at 70 rpm, leading to the formation of embryoid bodies (EBs). On day 2, the medium was changed to E6 medium (containing only hVEGF, hBMP4 and hSCF). On day 4 after mesoderm priming I, supernatant was discarded and 3 mL of mesoderm priming II medium (E6 medium supplemented with 50 ng/mL hVEGF and hBMP4, 20 ng/mL hSCF and 25 ng/mL hIL-3) was added and shaking speed was increased to 85 rpm. Mesoderm priming medium II was refreshed at day 7 before macrophage differentiation (hematopoietic differentiation) was started at day 10. For this, the medium was removed and the EBs were transferred to a 6-well tissue culture plate using 2 mL of differentiation medium (X-VIVO containing 1% (v/v) P/S and L-Glu, 0.1% (v/v) β-Mercaptoethanol supplemented with 25 ng/mL hIL-3 and 50 ng/mL hM-CSF. Macrophage production started from day 4 onwards and macrophages could be continuously harvested 1-2x per week.

#### Human iPSC-derived sensory neurons

iPSC cells (line LVPC118) were obtained from Dr. S. Viville, Strasbourg (Jung et al., 2014) and differentiated as previously described (Muller et al., 2018). Briefly, LVPC118 iPSC were seeded (day 0) into cell culture flasks coated with Geltrex (Thermofisher) at a density of 90,000 cells/cm2. Until day 10, LVPC118 were cultured in Basal medium, consisting of 1:1 (vol:vol) DMEM-F12 (Lonza) : NeuroBasal medium supplemented with 2% B-27, 1% N2, 1% MEM non-essential amino-acid (Thermofisher), 1% L-alanyl-L-glutamine, 0.1% trace element A, 0.1% trace element B, 0.1% trace element C (Corning), 50 μg/mL gentamycin (Lonza), 5 μM Y-27632, 0.01 mM β-mercaptoethanol (Thermofisher) and 50 μg/mL ascorbic acid (Sigma). From day 0 to day 4, Basal medium was supplemented with 40 μM SB431542, 0.2 μM LDN193189 and 3 μM CHIR99021 (Sigma). From day 5 to day 6, Basal medium was supplemented 3 μM CHIR99021. From day 9 to day 10, Basal medium was supplemented with 10 μM DAPT (Selleckchem). At day 11, neuron precursor cells (NPCs) were harvested using Accutase and cryopreserved for further use. NPCs were thawed and seeded in poly-D-lysin precoated wells with DMEM F12 and Neurobasal medium (1:1) containing N2 (Gibco, 100x), B27 (Gibco, 50x), 10 µM DAPT (Selleckchem), 50 µg/ml ascorbic acid (Sigma), 5 µM Y-27632 dihydrochloride (Tocris), 10 ng/ml human GDNF (Peprotech), 20 ng/ml human BDNF (Peprotech) and 0.5 % Streptomycin and Penicillin (Pan Biotech). Medium was changed the next day. From the third day on, DAPT was replaced by 10 ng/ml human NGF and medium was changed every other day.

### Cocultures

For mouse cell cultures, BMDM were added at a density of 25,000 cells/cm^2^ to DRG cultures (10,000 cells/cm^2^) and cocultured for 5 days. As a control, BMDM in monocultures were cultured in the same medium. In all approaches, 20 ng/ml M-CSF was added to the cultures. Medium was changed every second day. For experiments with conditioned medium, medium was taken from cocultures after two and four days. Medium was centrifuged and the supernatant was mixed 1:1 with fresh media for DRG cells, 20 ng/ml M-CSF and 10 ng/ml NGF. Conditioned medium was then added to BMDM on day two and four of monoculture. For transwell approaches, 50,000 BMDM were seeded into 24-well cell culture inserts with a PET membrane and 0.4 µm pore size (Carl Roth) were used which were placed on wells containing DRG cells. DRG medium with 20 ng/ml M-CSF and 10 ng/ml NGF was added and changed every second day. For human cocultures, iMacs were added at a density of 25,000 cells/cm^2^ to iPSC-derived sensory neurons (100,000 cells/cm^2^) at day 7 of differentiation and cocultured for 5 days. For cocultures, medium for sensory neurons containing NGF was used and Y-27632 dihydrochloride was replaced with 50 ng/ml human M-CSF. For control, iMacs were cultured alone in the same medium. Medium was changed every second day.

#### *In vitro* treatments

BMDM as well as human iMacs were treated with different concentrations (0.2 - 10 ng) of recombinant human TGF-β1 (Peprotech), reconstituted according to manufacturer’s instructions. For the inhibition of TGF-β signaling, 1 µM of TGF-β RI Kinase Inhibitor VIII SB-525334 (Cayman chemicals) was used. To inhibit integrin αVβ5, 5 µM Cilengitide (Tocris) was added to the cultures. For all treatments, cells were cultured with medium used for neuron cultures with additional 20 ng/ml murine M-CSF for mouse or 50 ng/ml human M-CSF for human macrophages. For tamoxifen inductions, 2mM OH-TAM in ethanol or ethanol alone was added to the culture, medium was exchanged after 3 days and cells were sorted based on YFP expression on day five.

### Flow cytometry and cell sorting

To block FcγII/III receptors, cells were incubated with anti-CD16/32 antibody (eBioscience) for 5min at 4°C in FACS buffer (PBS, 2 %FCS, 2 mM EDTA). Subsequently, the respective antibodies were added to a mastermix in FACS buffer for surface staining and incubation for 30 min at 4°C. To control for cell viability, DAPI or fixable viability dye^TM^ (eBioscience) were used according to manufacturer’s instructions. For analysis, a 3-laser flow cytometer (Gallios™, Beckman Coulter) was used and data were processed with the Kaluza software (v2.1, Beckman Coulter) or FlowJo (v10, LLC). Dermal cells and cells from cocultures were purified by cell sorting with a MoFlo Astrios AQ or a CytoFlex SRT (Beckman Coulter).

### Antibodies used for flow cytometry

The following anti-mouse antibodies were used for surface staining: CD3ε (145-2C11, BioLegend), CD11b (M1/70, Invitrogen), CD11c (HL3, BD), CD16/32 (93, Biolegend), CD45 (30-F11, Invitrogen), CD45.1 (A20, Invitrogen), CD45.2 (104, BioLegend), CD64 (X54-5/7.1, BioLegend), CD206 (C068C2, BioLegend), β3-tubulin (TuJ-1, R&D), CX3CR1 (SA011F11, BioLegend), F4/80 (Cl:A3-1, Thermo Fisher Scientific), Ly6C (HK1.4, BioLegend), Ly6G (1A8, BioLegend), MHC II (AF6-120.1, Invitrogen), NK1.1 (PK136, BioLegend), Siglec-F (S17007L, BioLegend), and LAP (TGF-β1) (TW7-16B4, BioLegend). The following anti-human antibodies were used for staining: CD11b (ICRF44, Invitrogen), CD14 (M5E2, BD Pharmingen), CD64 (10.1, BioLegend), CD206 (19.2, Invitrogen), CX3CR1 (2A9-1, BioLegend), and Neurofilament M (Poly28410, BioLegend).

### ELISA

For ELISA, cells were cultured in DRG medium without serum in 48-well plates. Medium was changed on second day of culture and collected on day five for analysis. TGF-β1 ELISA (RandD Systems, DY1679) was performed according to manufacturer’s instructions.

### RNA extraction and qRT-PCR

For RNA extraction, cell populations were sorted in RLT buffer containing 1 % β-mercaptoethanol. RNA was extracted with the Extractme total RNA micro spin kit or RNeasy Micro Kit Plus (Qiagen) according to manufacturer’s instructions. cDNA synthesis was performed by using SuperScript^TM^ IV VILO mix (Thermo Fisher). For qRT-PCR, ABsolute qPCR SYBR Green was used (Thermo Fisher Scientific) and samples were analyzed with a LightCycler 480 (Roche).

### RNAseq

Library preparation and RNAseq for dermal macrophages were performed at the Genomics Core Facility “KFB -Center of Excellence for Fluorescent Bioanalytics” (University of Regensburg, Regensburg, Germany; www.kfb-regensburg.de). The SMARTer Ultra Low Input RNA Kit for Sequencing v4 (Clontech Laboratories, Inc., Mountain View, CA, USA) was used to generate first strand cDNA from 250 pg total-RNA. Double stranded cDNA was amplified by LD PCR (13 cycles) and purified via magnetic bead clean-up. Library preparation was carried out as described in the Illumina Nextera XT Sample Preparation Guide (Illumina, Inc., San Diego, CA, USA). 150 pg of input cDNA were tagmented (tagged and fragmented) by the Nextera XT transposome. The products were purified and amplified via a limited-cycle PCR program to generate multiplexed sequencing libraries. For the PCR step 1:5 dilutions of index 1 (i7) and index 2 (i5) primers were used. The libraries were quantified using the KAPA Library Quantification Kit - Illumina/ABI Prism User Guide (Roche Sequencing Solutions, Inc., Pleasanton, CA, USA). Equimolar amounts of each library were sequenced on a NextSeq 500 instrument controlled by the NextSeq Control Software (NCS) v2.2.0, using two 75 Cycles High Output Kits with the dual index, single-read (SR) run parameters. Image analysis and base calling were done by the Real Time Analysis Software (RTA) v2.4.11. The resulting .bcl files were converted into .fastq files with the bcl2fastq v2.18 software. Trimmed (Cutadapt) raw reads were aligned to the mm10 murine reference genome with STAR (Version 2.7.2b) (Dobin et al., 2013) and quantified with feature counts (Liao et al., 2014).

RNA-sequencing and raw data processing for cell culture samples was performed by CeGaT (Tübingen, Germany). Libraries were prepared using 10 ng RNA and the SMART-Seq Stranded Kit (Takara). Sequencing was performed with 2x 100 bp read length and 50 million clusters/probe on NovaSeq 6000 (Illumina). Demultiplexing of the sequencing reads was performed with Illumina bcl2fastq (2.20). Trimmed raw reads were aligned to mm10 using STAR (version 2.7.3). Differential expression analysis between groups was performed with DESeq2 (version 1.24.0) (Love et al., 2014) in R (version 3.6.1) (R Core Team 2015). Data were analyzed with the RStudio software in order to perform principal component analysis between the samples and to visualize expression patterns in volcano plots. Heatmaps were generated with Morpheus (https://software.broadinstitute.org/morpheus/). Gene ontology enrichment analysis on upregulated DEGs was performed with Panther 18.0 (Thomas et al., 2022). The fgsea package on the Galaxy Freiburg platform (Afgan et al., 2022) was used for gene set enrichment analysis (GSEA). At indicated conditions, genes were preranked by log2 fold change and enrichment in the “TGF_BETA_SIGNALING_PATHWAY” was determined using a gene set merged from curated canonical pathway gene sets (M2, Molecular Signatures Database, https://www.gsea-msigdb.org/) including TGF-β related gene sets of Reactome, BioCarta and WikiPathways databases.

### Single cell RNA-seq reanalysis

Public data from mouse DRG in healthy conditions (Sharma et al., 2020) and human DRG (Tavares-Ferreira et al., 2022) were reanalyzed. The datasets were downloaded and the RDS file was imported into R (R Core Team, 2023) environment version v4.3.1 and Seurat v4.3.0 (Hao et al., 2021). For mouse DRG data, the pre-processing step was performed by filtering genes expressed in at least 300 cells and outlier cells based on three metrics (nCountRNA > 50,000 and nFeature_RNA > 6,500 and mitochondrial percentage expression > 5) were filtered out. The percentage of mitochondrial genes was regressed out using the SCTransform function and Principal Component Analysis (PCA) was performed using the top 45 PCs for dimension reduction. For pre-processing of human DRG data, cells based on three metrics (nCountRNA > 30,000 and nFeature_RNA > 6,000 and mitochondrial percentage expression > 10) were filtered out. The percentage of mitochondrial genes was regressed out in the SCTransform function and Principal Component Analysis (PCA) & dimension reduction was performed using the top 50 PCs. In both datasets, t-distributed stochastic neighbor embedding (t-SNE) was used for dimension reduction and clusters were identified using the author’s annotations. Visualization of genes illustrating expression levels was performed using R/Seurat commands (DimPlot, FeaturePlot, and DotPlot) with ggplot2 v3.4.2 and scCustomize v1.1.1 R packages.

### Histology

For ear skin whole mounts, mouse ears were split into dorsal and ventral parts, fixed overnight at 4°C in 1% paraformaldehyde (Electron Microscopy Sciences) and stained with primary and secondary antibodies diluted in washing buffer consisting of 1xPBS, 1%BSA and 0.025% (v/v) Triton X-100 (Sigma). For imaging of cell cultures, cells were fixed for 15 min with 4% Paraformaldehyde and stained with primary and secondary antibodies diluted in staining buffer containing 1x PBS, 10% normal goat serum (abcam), 0.05% Tween20 (Sigma-Aldrich).

For the detection of basement membranes, nerves and macrophages, the following antibodies were used: anti-collagen IV (ab19808, abcam), anti-beta-III tubulin NorthernLightsTM NL-637 (TuJ-1, R&D Systems), anti-F4/80 PE (BM8, Invitrogen), anti-F4/80 eFluor®660 (BM8, Invitrogen), anti-CD206 PE (19.2, Invitrogen). Secondary antibodies included goat anti-rabbit AF633, AF546 and AF488 (Invitrogen). Yellow fluorescent protein (YFP) was enhanced with anti-GFP DyLight488 antibody (600-141-215, Rockland). Confocal microscopy on skin whole mounts was performed with a LSM 880 or LSM980 confocal microscope equipped with a 20×/0.8 NA Plan-Apochromat objective (Carl Zeiss Microimaging) using 1-µm optical slices. Images are displayed as maximum-intensity projections of 10-200µm thick z-stacks. For three-dimensional reconstructions, 20-50µm volume were recorded in 0.5-1µm sections (2048×2048 pixel resolution) and analyzed with the IMARIS software (Bitplane, Switzerland). For evaluation of axon sprouting, 3mm x 3mm tiled images were recorded around the center of the ear punch with 140µm Z-stacks. Tiles and Z-stacks were merged and the area of β3-tubulin positive staining was evaluated using Image J (1.47v, NIH, USA). Images were taken with the same parameters on the same experimental day for each independent experiment and the color threshold was set according to controls of each group and applied to all other samples to ensure comparability.

### Live-cell imaging

For time-lapse live cell imaging, cells were labelled using cell tracker orange CMRA or cell tracker green (both ThermoFisher Scientific). Macrophages and neurons were stained separately before start of the coculture for 30 min in DMEM/F12 medium at 37°C and washed with medium. Cocultures were seeded in in 8-well µ-slides (ibidi, Germany). Time-lapse imaging was performed with a LSM 710 or CD7 LSM 900 microscope equipped with a 20x/0.8 NA Plan-Apochromat objective (Carl Zeiss Microimaging) and data was processed in ZEN (Zeiss, Germany).

### Oligonucleotide sequences for qRT-PCR

Mouse Oligonucleotides for qRT-PCR included the following sequences (5‘-3‘):

- *Gapdh* (for: ACTCCACTCACGGCAAATTC, rev: TCTCCATGGTGGTGAAGACA)
- *Il1b* (for: TTCAGGCAGGCAGTATCACTC, rev: GAAGGTCCACGGGAAAGACAC)
- *Axl* (for: TGAGCCAACCGTGGAAAGAG, rev: AGGCCACCTTATGCCGATCTA)
- *Plxna4* (for: GCAAGGAAGCTTTTGCCGAG, rev: TGAGTCCTTTCTCCACACGC)
- *Mrc1* (for: GGCTGATTACGAGCAGTGGA, rev: ATGCCAGGGTCACCTTTCAG)
- *Cx3cr1* (for: AGGACACAGCCAGACAAG, rev: TCAGGGGAGAAAGCAAG)
- *Ramp3* (for: GTGAGTGTGCCCAGGTATGC, rev: CGACAGGTTGCACCACTTC)
- *Ramp1* (for: CCATCTCTTCATGGTCACTGC, rev: AGCGTCTTCCCAATAGTCTCC)
- *Fcrls* (for: CTTCTGGTCTTCGCTCCTGTC, rev: ATGGTGTAGCTTGAAGCACTG)
- *Pmepa1 (*for: TGGAGTTCGTGCAAATCGTG, rev: TCCGAGGACAGTCCATCGTC)
- *F11r* (for: GTGCTTGTACCTCCATCCAAGCCG, rev: GAGGGTGGGGAACCATCATGCTC)
- *Mmp14* (for: AGGCTGATTTGGCAACCATGA, rev: CCCACCTTAGGGGTGTAATTCTG)
- *Itgb5* (for: GCTGCTGTCTGCAAGCACAA, rev: AAGCAAGGCAAGCGATGGA)
- *Zeb1* (for: TGAGCACACAGGTAAGAGGCC, rev: GGCTTTTCCCCAGAGTGCA
- *Gcnt2* (for: TCAGCCTATCTCTCATTGCTGC, rev: CGCTGTCCGCTGTATAAAAACT)
- *Smad7* (for: AAGTGTTCAGGTGGCCGGATCTCAG, rev: ACAGCATCTGGACAGCCTGCAGTTG

## QUANTIFICATION AND STATISTICAL ANALYSIS

Statistical analysis was performed with Prism 9.0 (GraphPad). For comparison of two groups, unpaired two-tailed student’s *t*-test was applied unless stated otherwise. For multiple comparisons, one-way ANOVAs were performed, followed by Tukey’s or Dunnett’s multiple comparison tests. Differences were considered statistically significant if p-values were ≤ 0.05. (ns, not significant; *p<0.05; **p<0.01; ***p<0.001, p****<0.0001). Data are represented as mean±SEM if not stated otherwise. Sample sizes and experimental repeats are indicated in the figure legends.

## SUPPLEMENTAL INFORMATION

### MOVIES

**Movie S1: Directed movement of macrophages in coculture with DRG neurons**

Images of cocultures were recorded after three days of culture with 20 min intervals for 17h. Macs were labeled with cell tracker orange prior to coculturing. Scale bar: 20µm.

**Movie S2: iMacs migrating along iPSC-derived sensory neurons in cocultures**

Images of coculture were recorded after three days of culture with 20 min intervals for 20h. iMacs were labeled with cell tracker orange and iPSC-derived sensory neurons with cell tracker green prior to co-culturing. Scale bar: 50µm.

---

**Figure S1.**
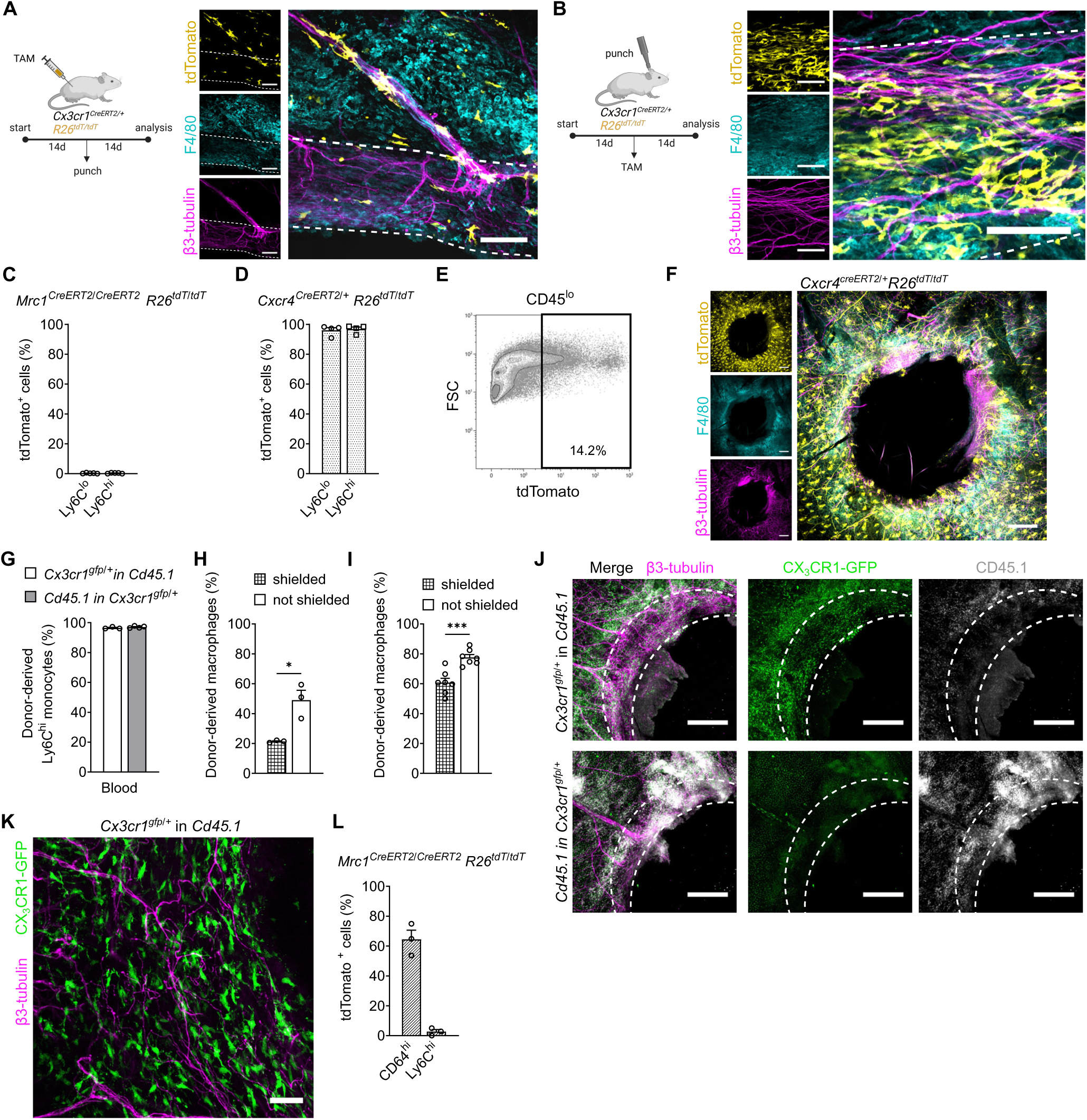
(A) Experimental setup and images showing lack of tdTomato labelling in the regeneration ring (dotted lines) of ear punches of *Cx3cr1^creERT2/+^ R26^tdT/tdT^* mice, induced prior to injury. Dashed lines indicate regeneration rings at injury center (n=4). Scale bar: 100µm. (B) Experimental setup and images showing tdTomato labelling in macrophages in the regeneration ring of ear punches of *Cx3cr1^creERT2/+^ R26^tdT/tdT^* mice, induced post injury (n=4). Scale bar: 100µm. (C) Quantification of tdTomato expression in peripheral blood monocytes of *Mrc1^creERT2/creERT2^ R26^tdT/tdT^* mice, measured in flow cytometry (n=5). (D) Quantification of tdTomato expression in peripheral blood monocytes of *Cxcr4^creERT2/+^ R26^tdT/tdT^* mice, measured in flow cytometry (n=5). (E) TdTomato expression in CD45^lo^ skin cells in *Cxcr4^creERT2/+^ R26^tdT/tdT^* mice, measured in flow cytometry. (F) Whole mount image depicting ear punch in *Cxcr4^creERT2/+^ R26^tdT/tdT^* mouse, 14d post injury (n=5). Scale bar: 500µm. (G) Quantification of donor-derived Ly6C^hi^ monocytes in peripheral blood of chimeric mice. (H) Donor-derived CD64^+^ dermal macrophages in shielded vs. non-shielded ears in homeostasis, quantified in flow cytometry (n=3 per group). (I) Donor-derived CD64^+^ dermal macrophages in shielded vs. non-shielded ears 14 days post punch, quantified in flow cytometry (n=7 per group). (J) Whole mount images depicting CX_3_CR1^high^ macrophages in ear punches in mice in which either donor (upper panel) or host (downer panel) cells were transgenic for *Cx3cr1^gfp/+^*(n=3-4 per group). Scale bar: 500µm. (K) Whole mount image depicting CX_3_CR1^high^ macrophages in ear punches in *Cd45.1* mice transplanted with *Cx3cr1^gfp/+^* bone marrow (n=3). Scale bar: 500µm. (L) Quantification of tdTomato^+^ cells in ear skin of *Mrc1^creERT2/creERT2^ R26^tdT/tdT^* mice, punched and induced with TAM 14 days prior to analysis, quantified via flow cytometry (n=3). Data are mean±SEM. *p<0.05, ***p<0.001, ns, not significant (two-tailed unpaired *t*-test). See also Figure 1.

**Figure S2.**
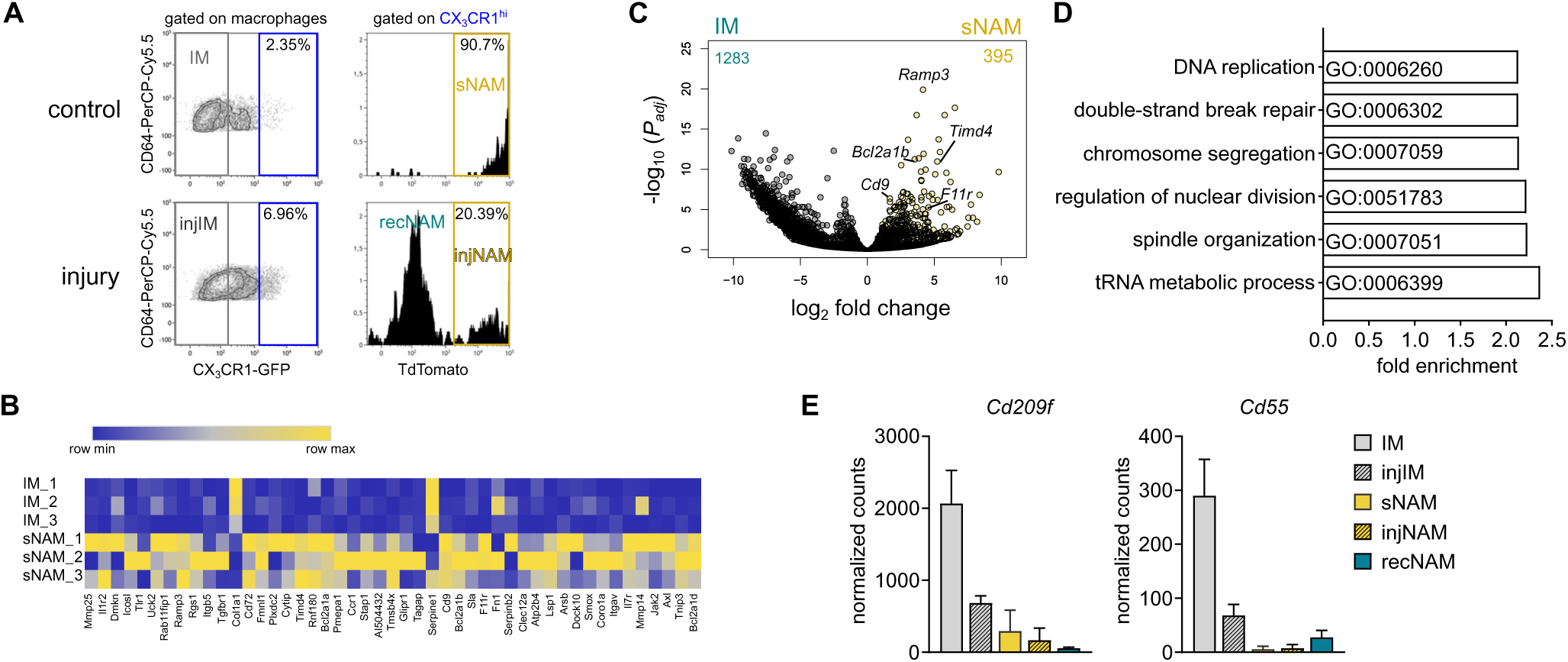
(A) Gating and sorting strategy for sequenced samples of Figure 2. (B) Heatmap for top 50 sNAM signature genes in IM and sNAM in steady state. (C) Volcano plots of differentially expressed genes in IM vs. sNAM. Colored dots represent a log2-transformed fold change >1 and p≤ 0.05 (Padj = two-tailed Benjamini-Hochberg test P value). (D) GO terms enriched in sNAM upon injury. Depicted are the highest enriched terms including at least 150 genes. FDR= False discovery rate (<0.05). (E) Normalized counts of selected IM genes, which are downregulated in sNAM. ata are mean±SEM. See also Figure 2.

**Figure S3.**
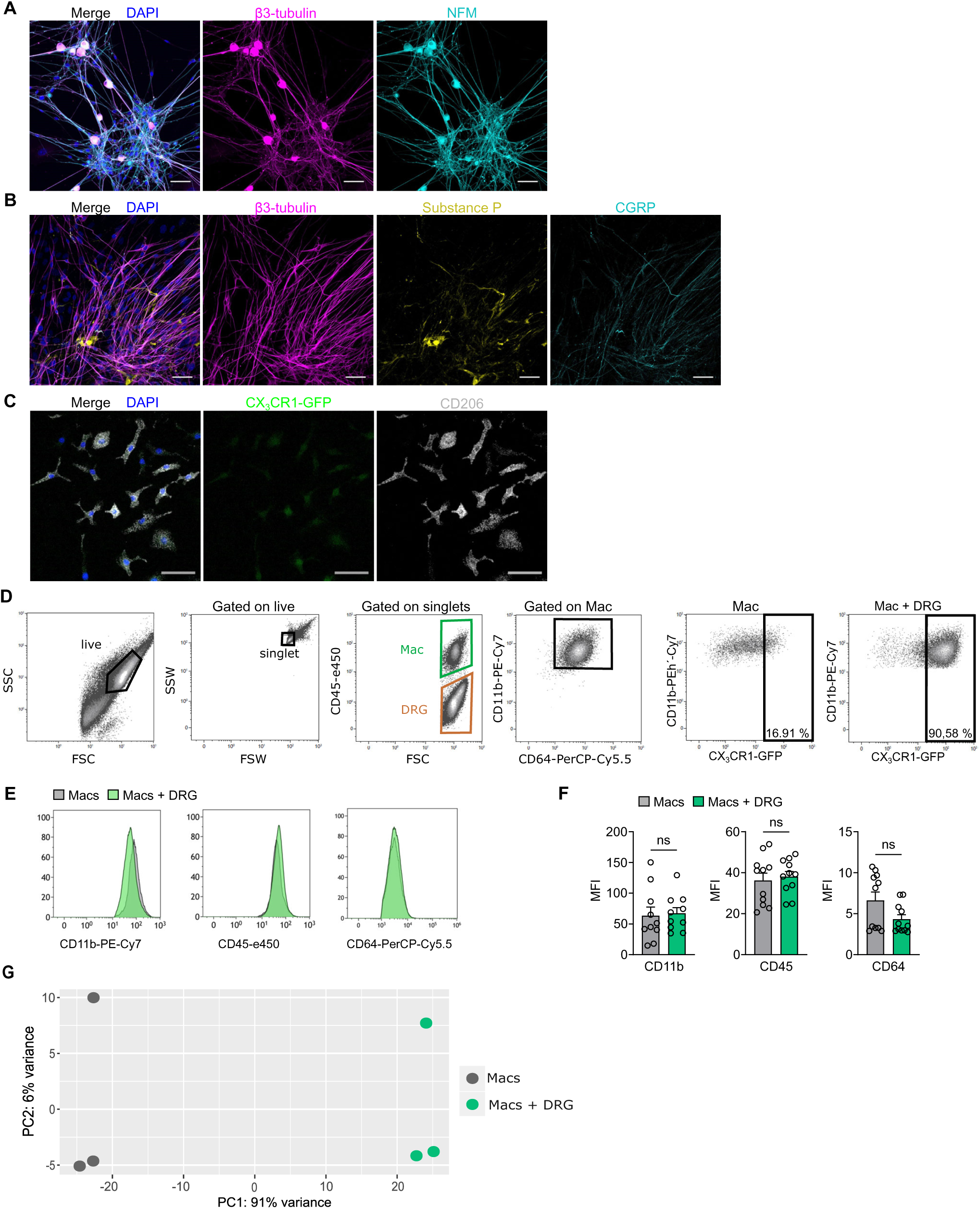
(A) NFM staining of sensory neurons after 12 days in culture. Scale bar: 50 μm (B) IF staining of neuropeptides expressed by sensory neurons after 12 days in culture. Scale bar: 50 μm (C) IF staining of CX3CR1-GFP Macs in monoculture. Scale bar: 50μm. (D) Gating strategy and CX3CR1-GFP expression in Macs cultured alone or with DRG neurons. (E) Histogram depicting CD45, CD11b and CD64 expression by Macs in cocultures. (F) Comparison of MFI of Mac surface markers measured by flow cytometry (n=11). (G) PCA analysis of Macs in mono- and coculture. Data are mean±SEM. ns= not significant (two-tailed unpaired *t*-test). See also Figure 3.

**Figure S4.**
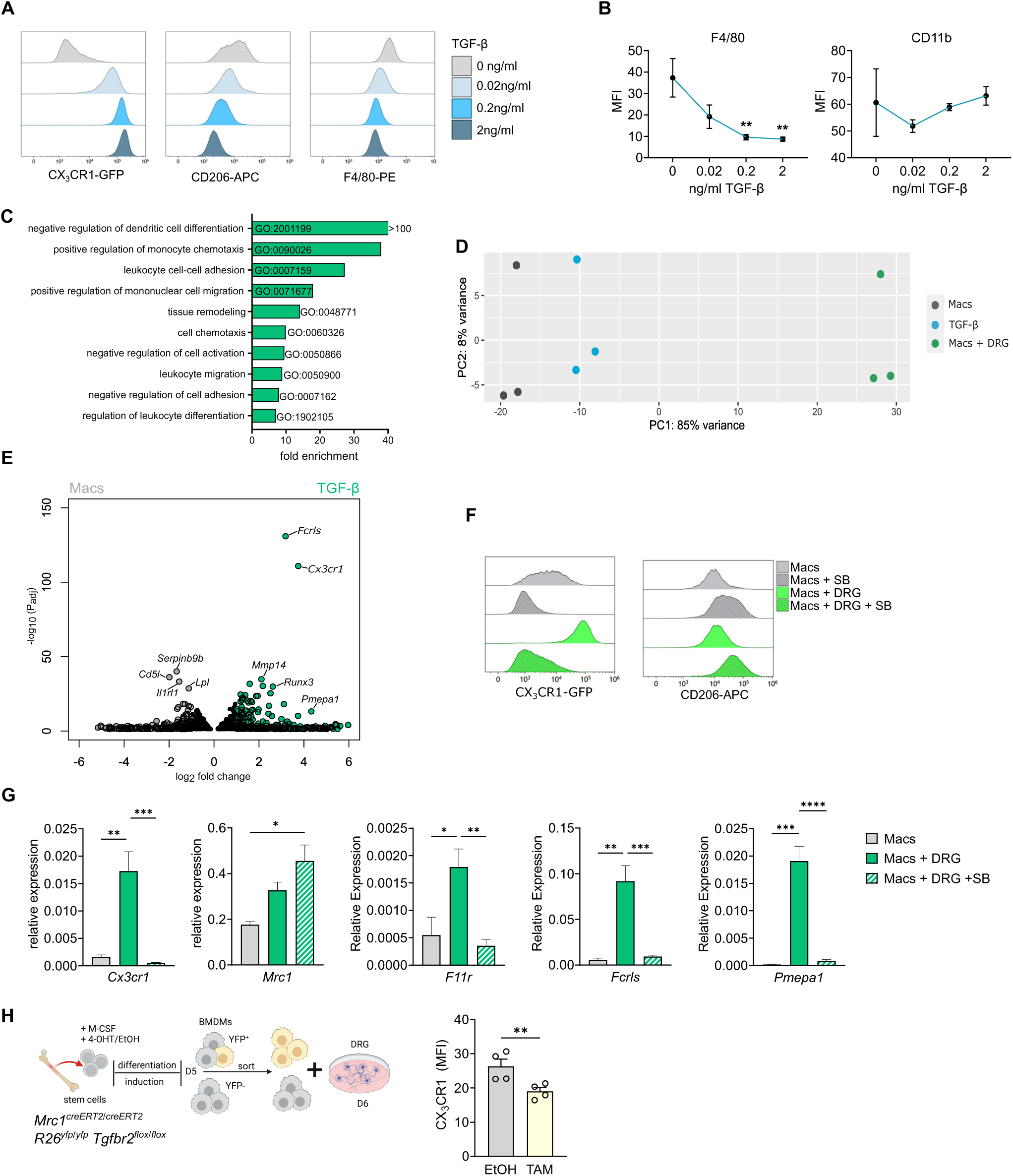
(A) Histograms of CX_3_CR1-GFP, CD206 and F4/80 expression in Macs treated with different concentrations (0.02-2 ng/ml) of TGF-β for 5 days, measured by flow cytometry. (B) MFI of CD11b and F4/80 in Macs cultivated for 5 days with different concentrations of recombinant TGF-β, quantified by flow cytometry (n=4). (C) GO slim biological process pathways enriched in Macs treated with 0.02ng TGF-β for 5 days. Visualization of terms with highest fold enrichment in genes with a base mean >100. (D) PCA analysis of Macs cultured either alone, with DRG or with 0.02 ng of TGF-β. (E) Volcano plot of differentially expressed genes in untreated Macs and treated Macs with 0.02 ng TGF-β for 5 days. Grey/green dots represent log2-transformed fold change >1. Padj = two-tailed Benjamini-Hochberg test P value. (F) Comparison of CX_3_CR1-GFP and CD206 expression in Macs in mono- and coculture with or without TGFBR1 inhibitor SB525334 (SB). (G) Relative gene expression of *Mrc1* and the depicted signature genes of Macs in monoculture, coculture and cocultures treated with TGFBR1 inhibitor SB525334 (SB) (n=3-5 per group). (H) Cocultures with Macs derived from *Mrc1^creERT2/creERT2^ R26^yfp/yfp^ Tgbfr2^flox/flox^* mice. Macs were treated with EtOH or OH-TAM during differentiation, YFP^-^ and YFP^+^ cells were sorted and added to DRG cultures. MFI of CX_3_CR1 was quantified in Macs by flow cytometry after coculturing for 5 days (n=4) (two-tailed paired *t*-test). Data are mean±SEM. *p<0.05, **p<0.01, ***p<0.001, ****p<0.0001, ns, not significant (one-way ANOVA, Tukey’s multiple comparison test if not stated otherwise). See also Figure 4.

**Figure S5.**
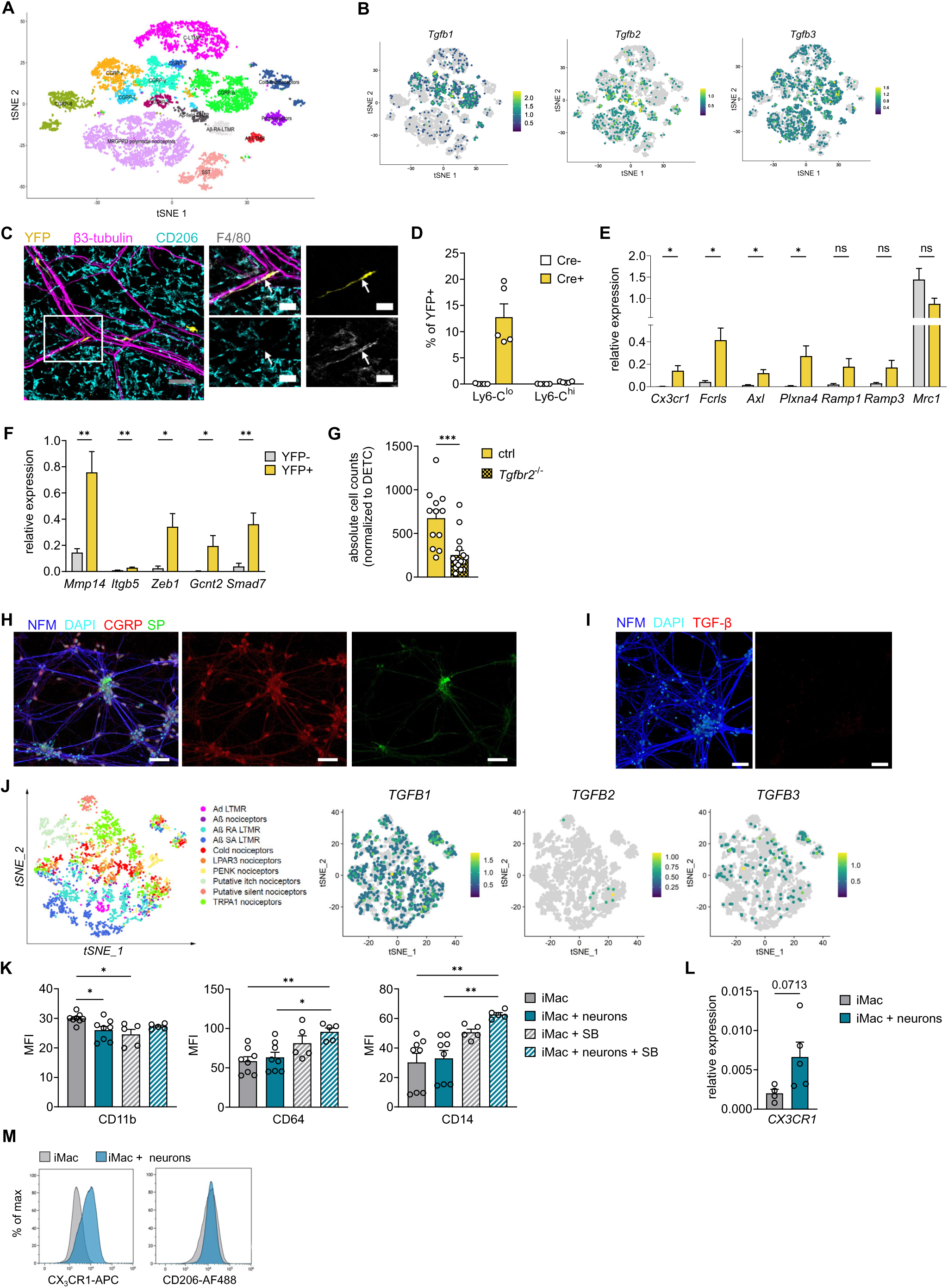
(A) tSNE map visualisations of scRNAseq data of mouse DRG (GSE19088, Sharma et al, 2020 Nature). (B) tSNE feature plots for *Tgfb1-3* expression from GSE19088. (C) Whole mount image depicting *Cx3cr1^creERT2/+^ R26-yfp^flox/flox^* ear skin, which were used as controls, induced with TAM 4 weeks prior to analysis. Scale bar: 50µm, zoom in: 20µm. (D) Recombination efficacy in circulating monocyte subsets in *Cx3cr1^creERT2/+^ R26-yfp^flox/flox^* 7 days post TAM administration. n=5. (E) Relative gene expression of sNAM signature genes in sorted YFP^-^ and ^+^ macrophages from *Cx3cr1^creERT2/+^ R26-yfp^flox/flox^* mice. n=8 per group, 3 independent experiments. (F) Relative gene expression of TGF-β related genes in sorted YFP^-^ and ^+^ macrophages from *Cx3cr1^creERT2/+^ R26-yfp^flox/flox^* mice. n=8 per group, 3 independent experiments. (G) Absolute counts of YFP^+^ macrophages in ear skin of *Cx3cr1^creERT2/+^ R26-yfp^flox/flox^Tgfbr2^flox/flox^* mice compared to controls. n=12-16 per group, 5 independent experiments (two-tailed unpaired *t*-test). (H) IF staining of neuropeptides in iPSC-derived sensory neuron cultures. Scale bar: 50 µm. (I) TGF-β staining in iPSC-derived sensory neurons cultured without iMacs. Scale bar: 50 µm. (J) tSNE map and feature plot of *TGFB1-3* expression of scRNAseq data of human DRG (Tavares-Ferreira et al, 2022). (K) MFI for depicted surface markers of iMacs in mono-and cocultures with or without TGFBR1 inhibitor SB525334 (SB) measured by flow cytometry (n=5-8 per group). (L) Relative gene expression of *CX3CR1* in iMacs sorted from monocultures or cocultures with iPSC-derived neurons (two-tailed unpaired *t*-test) (n=4). (M) CX_3_CR1 and CD206 expression of iMacs cultured alone or added during differentiation of iPSC-derived sensory neurons and cocultured for 12 days. Data are mean±SEM. *p<0.05, **p<0.01, ***p<0.001, ****p<0.0001 (one-way ANOVA, Tukey’s multiple comparison test if not stated otherwise). See also Figure 5.

**Figure S6.**
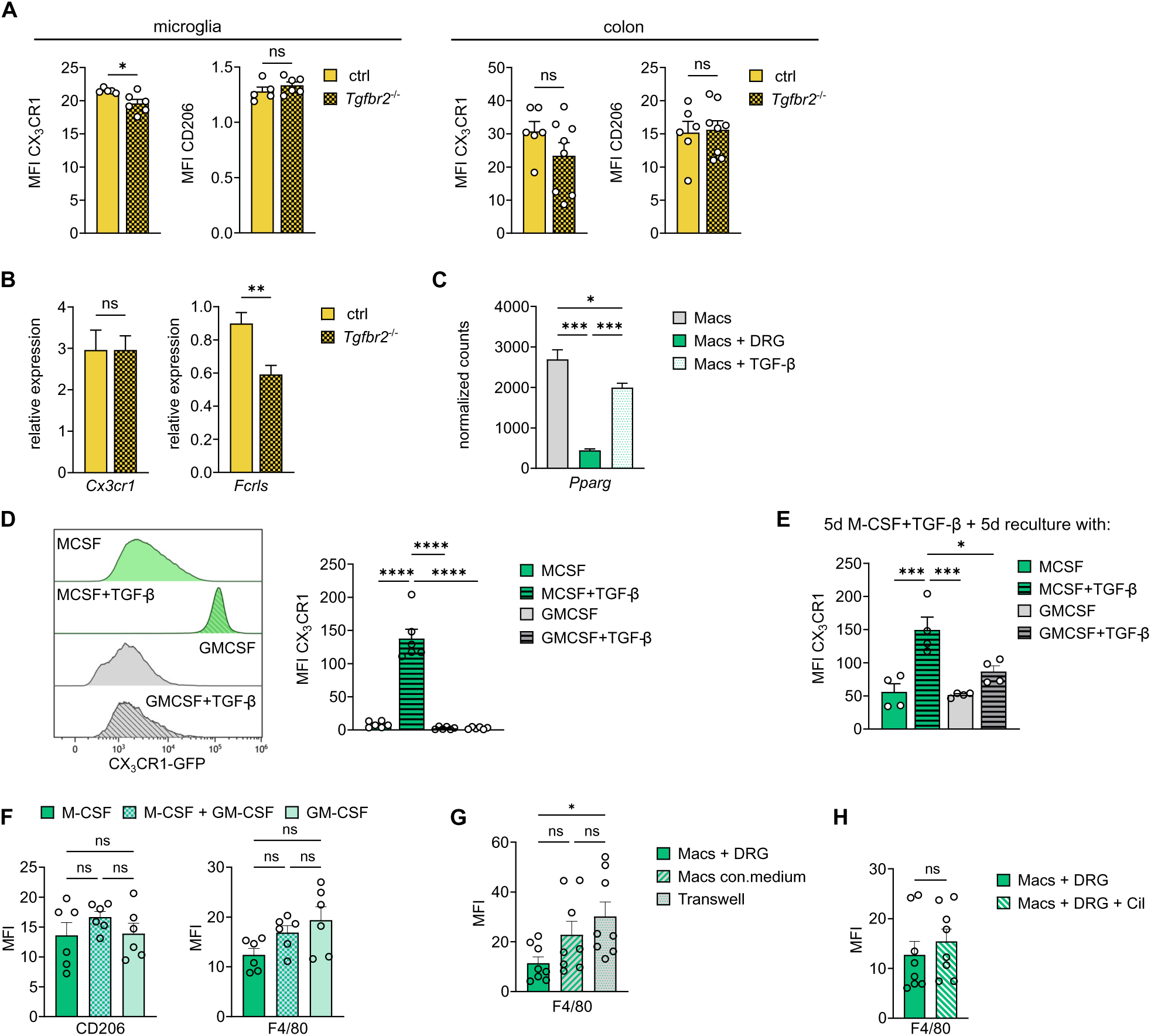
(A) MFI of CX_3_CR1 and CD206 in YFP^+^ microglia and colon macrophages 4 weeks post TAM (two-tailed unpaired *t*-test). (B) Relative expression of selected genes in microglia, sorted 4 weeks post TAM (two-tailed unpaired *t*-test) (n=5-6 per group). (C) Normalized counts of *Pparg* in Macs, Macs in coculture with DRG and Macs treated with 0.02 ng/ml TGF-β. (D) CX_3_CR1 expression by Macs differentiated with M-CSF or GM-CSF ± 2ng/ml TGF-β for 10 days. (E) CX_3_CR1 expression by Macs differentiated with M-CSF+2ng/ml TGF-β for 5 days and then recultured with M-CSF or GM-CSF ± 2ng/ml TGF-β for another 5 days (n=4). (F) Comparison of CD206 and F4/80 expression in Macs cocultured with DRG neurons and either with M-CSF, GM-CSF and M-CSF or with GM-CSF only, measured by flow cytometry (n=6). (G) MFI for F4/80 in Macs either cocultured with DRG, treated with conditioned medium (con. medium) from cocultures or cultured in transwells with DRGs (n=8). (H) MFI for F4/80 in Macs in coculture with DRGs treated with ανβ5 inhibitor cilengitide (Cil) (two-tailed unpaired *t*-test) (n=8).

**Figure S7.**
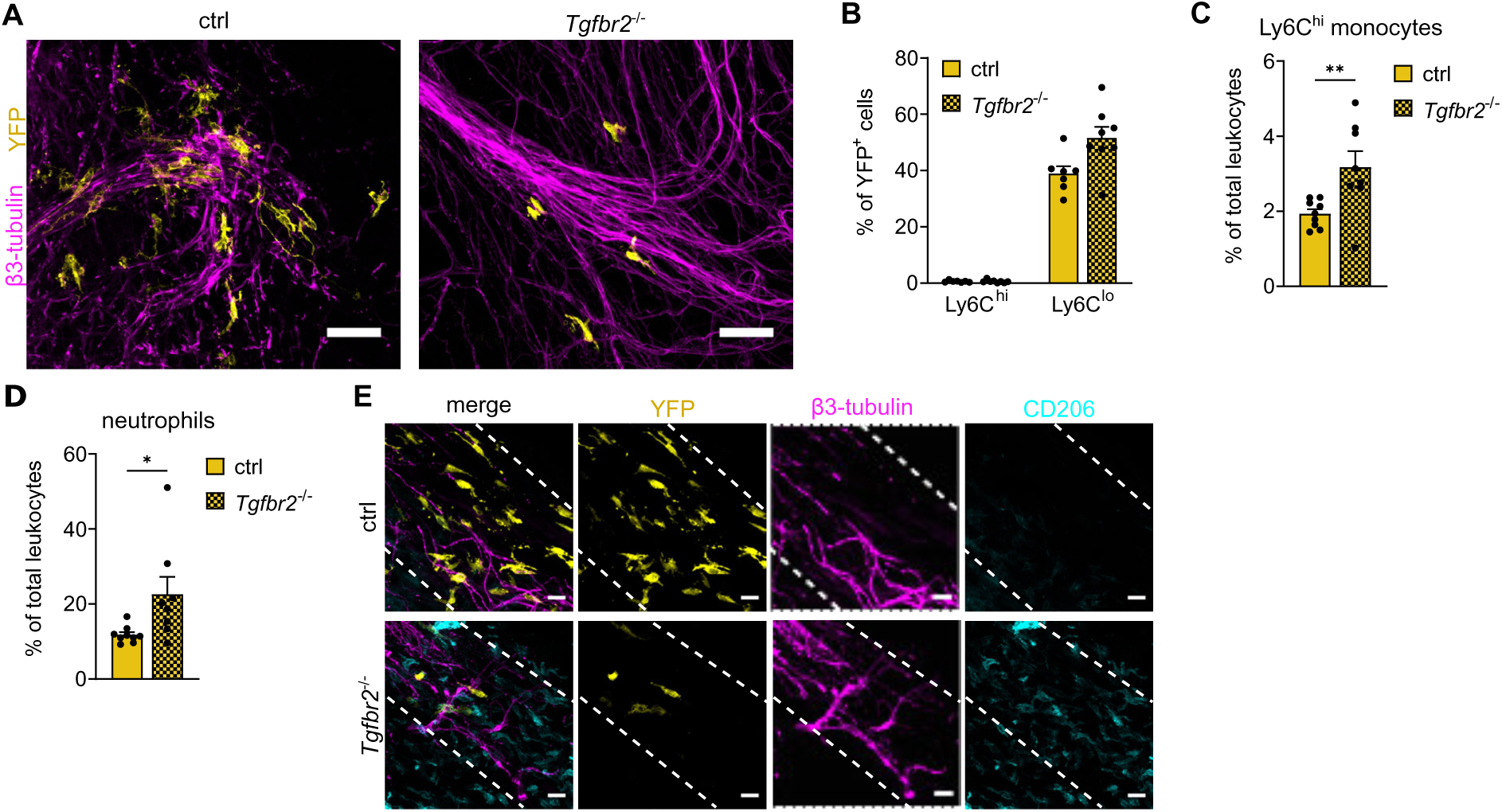
(A) YFP^+^ macrophages at injured nerve endings (related to Figure 7A). Scale bar: 50µm. (B) Proportion of YFP^+^ cells among Ly6C^hi^ and ^lo^ blood monocytes. n=7-8 per group. (C) Quantification of Ly6C^hi^ monocytes in peripheral blood of mice described in Figure 7D. n=8-9 per group. (D) Quantification of Ly6G^+^ neutrophils in peripheral blood via flow cytometry. (E) YFP^+^ macrophages in regeneration rings of *Cx3cr1^creERT2/+^ R26-yfp^flox/flox^Tgfbr2^flox/flox^* and controls upon continuous TAM administration. Data are mean±SEM. *p<0.05 (two-tailed unpaired *t*-test). See also Figure 7.

